# A deep-learning strategy to identify cell types across species from high-density extracellular recordings

**DOI:** 10.1101/2024.01.30.577845

**Authors:** Maxime Beau, David J. Herzfeld, Francisco Naveros, Marie E. Hemelt, Federico D’Agostino, Marlies Oostland, Alvaro Sánchez-López, Young Yoon Chung, Michael Maibach, Stephen Kyranakis, Hannah N. Stabb, M. Gabriela Martínez Lopera, Agoston Lajko, Marie Zedler, Shogo Ohmae, Nathan J. Hall, Beverley A. Clark, Dana Cohen, Stephen G. Lisberger, Dimitar Kostadinov, Court Hull, Michael Häusser, Javier F. Medina

## Abstract

High-density probes allow electrophysiological recordings from many neurons simultaneously across entire brain circuits but don’t reveal cell type. Here, we develop a strategy to identify cell types from extracellular recordings in awake animals, revealing the computational roles of neurons with distinct functional, molecular, and anatomical properties. We combine optogenetic activation and pharmacology using the cerebellum as a testbed to generate a curated ground-truth library of electrophysiological properties for Purkinje cells, molecular layer interneurons, Golgi cells, and mossy fibers. We train a semi-supervised deep-learning classifier that predicts cell types with greater than 95% accuracy based on waveform, discharge statistics, and layer of the recorded neuron. The classifier’s predictions agree with expert classification on recordings using different probes, in different laboratories, from functionally distinct cerebellar regions, and across animal species. Our classifier extends the power of modern dynamical systems analyses by revealing the unique contributions of simultaneously-recorded cell types during behavior.

---

The nervous system comprises many molecularly, anatomically, and physiologically defined cell types^1–6^. Powerful modern molecular techniques now have revealed multiple sub-types even within known anatomical cell classes^7–13^. Identification of cell type at multiple levels will be crucial to understand how the brain works and to develop selective, targeted therapeutics for brain dysfunction. Therefore, it is crucial to develop strategies to determine cell type and to cross-reference different formulations of cell type across levels of analysis^3,5,6,12,14,15^.

With the advent of high-density multi-contact recording probes^16,17^, it is now possible to record from hundreds of neurons simultaneously and characterize their activity during specific, quantified behaviors. Simultaneous large-scale electrophysiological recordings coupled with cell-type identification *in vivo* would facilitate characterization of circuit-level processing in the service of behavior. Yet, identification of cell type is a particularly difficult challenge for extracellular recording technologies that cannot access the transcriptional or anatomical properties of neurons^18^. Efforts to classify neurons based on specific features of their spike waveform and firing statistics have not proven robust across laboratories^19,20^. Moreover, optogenetic approaches to cell-type identification^21–24^ currently are routine only in mice and bring the technical challenges of (i) off-target expression of opsins^25^, (ii) disambiguating direct responses versus those due to recurrent connectivity within circuits^26^, and (iii) the ability to target only one or two cell types at a time in a given preparation^27^.

We assembled a collaboration of four laboratories with the single-minded goal of enabling cell-type identification solely from extracellular recordings in awake animals by developing a strategy that could scale across labs, probes, species, and in the future maybe across brain areas. We chose to pioneer the strategy in the cerebellar cortex, which provides key advantages. Specifically, the cerebellum has a crystalline architecture with well-defined neuronal connectivity and a small number of anatomically-defined cell types^1,28^ that are consistent across species^29,30^, allowing direct comparison of recordings in monkeys, mice and other species. The cerebellum has a range of neuron sizes from among the smallest and most densely packed (granule cells) to the largest (Purkinje cells), allowing us to test the resolution of our recording approaches. The cerebellum has many spontaneously firing neurons^31–33^, some with high spontaneous rates, allowing us to extract rigorous information about their electrophysiological properties. Genetically-defined mouse Cre-lines are available for all major cell types in the cerebellum^34–38^, allowing us to leverage optogenetic strategies for cell-type identification^21^. Finally, the cerebellum has a long history of neurophysiological recording^39^, allowing us to reference our measurements and automated cell-type classifications against hard-won human expertise. Strategies to solve the challenges of cell-type identification in such a testbed should provide a roadmap for application to other structures, including the cerebral cortex, the hippocampus, and the basal ganglia.

Our approach succeeded. We created a ground-truth library of identified cerebellar cell types recorded in unanesthetized mice by combining rigorous spike sorting and unit curation with identification through combined optogenetic activation and pharmacological synaptic blockade. We demonstrate that a semi-supervised deep-learning classifier accurately predicts cell type for the ground-truth library based on the waveform, discharge statistics, and anatomical layer of the recording. Importantly, the classifier identifies cell type with high confidence in a high fraction of expert-labeled cerebellar recordings from two different laboratories, in behaving mice and macaque monkeys. When used for modern dynamical systems analysis, our classification strategy enables new biological insights by revealing the distinct temporal dynamics of simultaneously recorded, identified cell types during complex behaviors in mice and monkeys. Our classifier can therefore be leveraged as a key tool for testing how specific cell types support neural computation and behavior.

## Results

### General approach

We start by creating a ground-truth library of extracellular recordings from neurons whose cell type is established unequivocally. The analysis and classification pipeline (Figure 1) begins with several data curation steps to ensure high-quality recordings and allow us to characterize with high confidence each neuron’s waveform, resting discharge properties, and anatomical location. We use well-characterized mouse Cre-lines to identify cell types by optogenetic activation of specific cell types in the presence of synaptic blockers. We then develop a semi-supervised deep-learning classifier with performance evaluated with leave-one-out cross-validation. Finally, we use the classifier to predict the cell types of an independent dataset of recordings made in mice and macaque monkeys and we compare the performance of the classifier against cell-type identification by human experts. Below, we develop the details of our strategy one step at a time.

**Figure 1:**
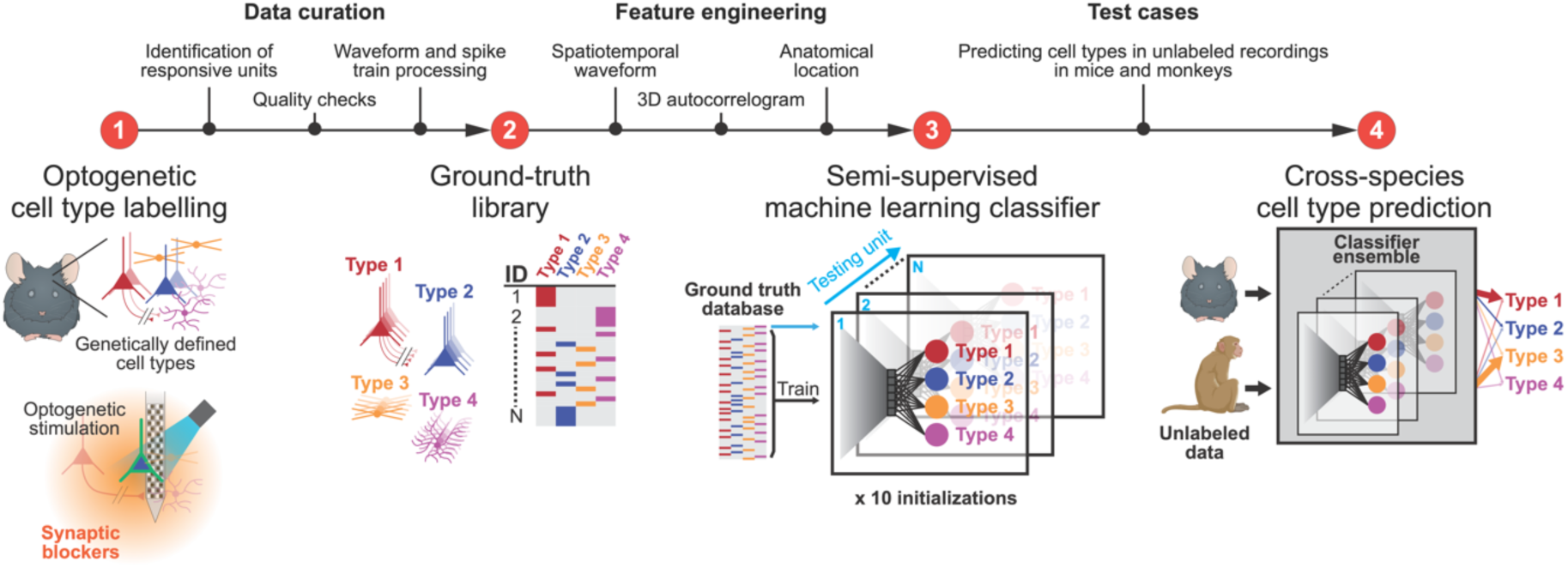
A strategy for cell type identification from extracellular recordings in neural circuits. The strategy comprises three steps: data acquisition and curation to build a ground-truth cell type library, selection of features from the ground-truth library to train a machine-learning based classifier, and tests of the classifier using additional datasets, including from other species. The first step is to create a ground-truth library of cell types based on optogenetic activation of genetically-defined neurons during electrophysiological recordings in awake mice. Neurons in the ground-truth library must be activated directly, as confirmed by a combination of synaptic blocker pharmacology and electrophysiological criteria, followed by careful data curation. The second step is to identify features in the dataset that can be used to train a semi-supervised deep-learning classifier. The third step is to test the generality of the classifier by asking it to predict cell types in independent datasets of expert-classified recordings from mice and monkeys.

### Multi-contact probe recordings and data curation

We develop and deploy the general strategy for cell-type classification (Figure 1) in the cerebellum, based on ground-truth recordings with Neuropixels probes in two laboratories (Häusser and Hull labs). In the cerebellar cortex, morphologically distinct cell types reside in different layers (Figure 2A). Purkinje cells comprise a monolayer and extend their planar dendrites through the molecular layer. Molecular layer interneurons reside across the extent of the molecular layer and include basket cells that innervate the Purkinje cell’s soma and stellate cells that innervate the Purkinje cell’s dendrites. The granule cell layer includes mossy fiber terminals, Golgi cells, and granule cells. Recent studies have identified other, less-common cell types in the different layers^37^, but we have elected to focus on the main cell types from the cerebellar circuit (Figure 2A).

**Figure 2:**
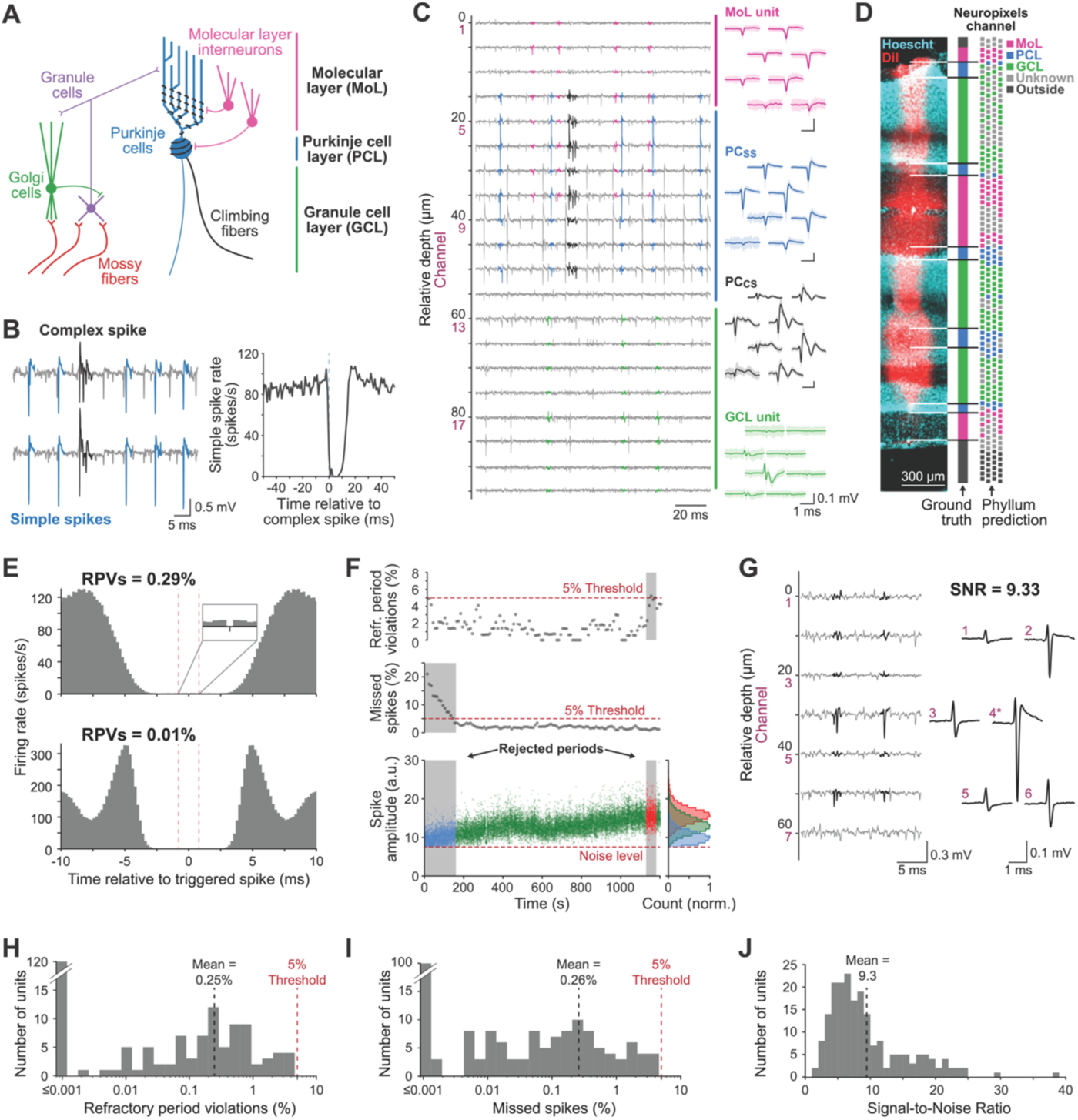
Curation of Neuropixels recordings in the mouse cerebellar cortex. A. Schematic diagram of the canonical cerebellar circuit. B. Traces on the left show example simple spikes (light blue) and complex spikes (black) in a Purkinje cell. Histogram on the right documents a complex-spike-triggered pause in simple spikes. C. Example recordings from many channels of a Neuropixels probe with magenta, blue, black, and green used to highlight a single unit recorded in the molecular layer, a Purkinje cell’s simple spikes, the same Purkinje cell’s complex spikes, and a unit recorded in the granule cell layer. D. Comparison of example histology labeled with DiI and Hoechst to show the excellent agreement of histological determination of layers and the layers predicted by Phyllum from the electrical recordings. Different colors on the Neuropixels schematic show: magenta, molecular layer; blue, Purkinje cell layer; green, granule cell layer; gray, unknown layer; black, outside cerebellar cortex. E. Autocorrelograms plotting a neuron’s firing rate as a function of time from one of its own trigger spikes for two neurons with very few refractory period violations (RPVs). Note that the spike counts in the autocorrelograms have been divided by the width of the bin so that the y-axis is in spikes/s. F. Analysis of quality of isolation as a function of time during a recording session. From top to bottom graphs show the percentage of refractory period violations, the estimated percentage of missed spikes, and spike amplitude. Horizontal dashed lines show thresholds for acceptance. Gray regions show periods that were rejected from analysis. Blue, green, and red symbols indicate spikes that came from intervals that had too many missed spikes, acceptable isolation, and too many refractory period violations. Marginal histograms on the right show the distribution of spike amplitudes to document clipping at the noise level in the blue histogram that would be cause for rejection of a time interval. G. Example recording traces and spatial footprint of a representative recording with a signal-to-noise ratio (SNR) of 9.33, with the waveforms numbered according to their channel. Asterisk (*) denotes the channel with the largest peak-to-trough amplitude, used to compute the SNR. H. Distribution of percentage of refractory period violations across neurons accepted to the ground-truth library. I. Distribution of estimates of percentage of spikes that were missed across neurons accepted to the ground-truth library. J. Distribution of signal-to-noise ratios on the channel with the largest-amplitude waveform across neurons accepted into the ground-truth library.

Purkinje cells are the one cell type in the cerebellum that allows ground-truth identification from its extracellular electrical signature. Purkinje cells show two types of action potentials (Figure 2B, left panel): “simple spikes” that fire at high rates and “complex spikes” driven directly by climbing fiber input^40–42^. Complex spikes occur only at ∼1 Hz, and trigger a characteristic 10-50 ms pause in simple spikes^43^. Thus, Purkinje cells can be identified unequivocally, and admitted into the ground-truth library, if they show a pause in a complex-spike-triggered histogram of simple-spike firing (Figure 2B, right panel).

Recordings with Neuropixels probes detect neural activity on many of the 384 channels and spike sorting yields many units including non-Purkinje cells. The magenta waveforms in Figure 2C arise from a neuron in the molecular layer that would be a candidate to be a molecular layer interneuron. The green waveforms come from a neuron recorded in the granule cell layer that could be a mossy fiber, a Golgi cell, or a granule cell. The blue and black waveforms are the simple spikes and complex spikes of an identified Purkinje cell.

Given that the soma of each cell type resides in one of the three different layers of the cerebellar cortex, the first step in our analysis pipeline was an objective procedure to identify the layer of each recording. The cerebellum is a foliated structure so that a single penetration with a Neuropixels probe usually records from neurons in multiple repetitions of each of the 3 layers of the cerebellar cortex. For example, the recording trajectory documented with DiI staining in Figure 2D crossed 3 molecular layers, 5 Purkinje cell layers, and 3 granule cell layers. We assigned each channel to a layer using Phyllum, a Phy plugin that analyzes recordings across the channels on a probe to infer the layer recorded by each channel (see ***Methods***).

The layer structure inferred by Phyllum agreed well with histological data based on simultaneous DiI and cell body staining (Figure 2D). We validated Phyllum across 21 histologically confirmed penetrations and found that its conclusions agree with the histology at 99, 95, and 98% of 776, 367, and 1140 recording sites respectively in the molecular, Purkinje cell, and granule cell layers.

The layer assignments from Phyllum were also consonant with the finding from single electrodes of (i) clear complex-spike activity and well-isolated simple spikes in the Purkinje cell layer, (ii) relative silence and abundant dendritic Purkinje cell complex spikes^44,45^ in the molecular layer, and (iii) a jungle of high-intensity activity with many units in the granule cell layer.

We next ensured that each unit we admitted for further analysis was a well-isolated single neuron with credible waveform and resting discharge properties, two of the three features we ultimately would use, along with layer, to classify units. We manually curated the output from Kilosort2 with Phy and subsequently performed automated quality checks to ensure the quality of isolation and the veracity of the waveforms and resting discharge statistics of neurons that would become part of our ground-truth library. We strove to ensure that we neither missed many spikes from the neuron under study nor included electrical artifacts or spikes from neighboring neurons.

● We analyzed the refractory periods from each isolated neuron to assess the level of contamination from other neurons or noise^46^. The examples of autocorrelograms in Figure 2E have vanishingly small numbers of refractory period violations and respectively represent the mean (0.25%) and median (0.01%) in our dataset. We rejected from the ground-truth library autocorrelograms with greater than 5% period violations (Figure 2F, red symbols and histogram). Almost all accepted neurons had fewer than 1% refractory period violations with a mean of 0.25% (Figure 2H).
● We estimated the number of missed spikes by fitting the spike amplitude distribution with a Gaussian function and quantifying the fraction of the area under the curve that was clipped at noise threshold^47,48^ (Figure 2F). We estimated that few spikes were missed if the distribution of spike amplitudes for the entire recording was continuous and not clipped at noise threshold. In Figure 2F, we estimated that more than 5% of spikes were missed in the first ∼150 s of the recording (blue symbols and histogram), causing us to exclude those intervals from further analysis. Among the recordings we accepted, the percentage of missed spikes averaged 0.26% and almost all neurons showed fewer than 1% missed spikes (Figure 2I).

The requirement for few violations of the refractory period and small numbers of missed spikes ensured that the units we accepted had high signal-to-noise ratios. The mean signal-to-noise ratio in our accepted sample, measured as the signal-to-noise ratio on the channel with the largest unit potential, was 9.3, almost identical to that of the example recording in Figure 2G. Over 90% of the neurons had signal-to-noise ratios larger than 4 (Figure 2J).

We identified and resolved two other issues that impaired consistent and reliable estimates of waveform. The *<u>first</u>* issue is related to an on-board hardware high-pass filter on Neuropixels probes. The filter distorts the shape of waveforms and therefore hinders comparison across recordings made when the filter was on versus off. We used the technical description of the analog filter in the Neuropixels documentation to apply an equivalent digital filter to data recorded with the filter disengaged or using other probes in monkeys (Supplementary Figure 1). The *<u>second</u>* issue concerns temporal alignment of individual spikes, which is unreliable in Kilosort’s output when the signal-to-noise ratio is low to medium or when a unit drifts across channels during a recording session. We resolve the alignment issue with an iterative procedure we called “drift-shift-matching” to minimize waveform distortion from the averaging of individual action potentials (see Supplementary Figure 1 and ***Methods***).

### Combination of optogenetics and pharmacology for ground-truth cell-type identification

For optogenetic activation, we combined genetic and viral approaches to cause expression of an opsin in a specific cell type, thereby allowing these neurons to be selectively activated by light and identified by photostimulation^21^. We added synaptic blockers to our experimental preparations to ensure that light activates an opsin-expressing neuron directly, and not indirectly by optogenetic activation of its pre-synaptic inputs. The standard criterion of short-latency activation by optogenetic stimulation^49^ (e.g., <10 ms) is inadequate on its own. We found ample examples of short-latency responses (some <5 ms) that disappeared with synaptic blockade^21^.

We introduced Neuropixels probes into the lateral cerebellar cortex or the vermis of mice expressing opsins (usually Channelrhodopsin-2, ChR2) in specific cell types and allowed the probe to settle at a location where we recorded activity across much of its length. As illustrated in Figure 3A, we performed experiments with optogenetic stimulation in multiple phases. In a *baseline phase*, we recorded spontaneous activity. In a *control phase*, we applied light externally to the cerebellum to activate opsins in the cell types that expressed them. We also introduced a tapered optic fiber that ran alongside the recording probe in some experiments, to deliver light in closer proximity to cells that expressed opsins. In an *infusion phase*, we continued to deliver light to the cerebellum while we added synaptic blockers (see ***Methods***) to the surface of the cerebellum. In the *blockade phase*, when the synaptic blockers had permeated well into the tissue, we assayed neurons for direct responses to optogenetic stimulation. The approach of applying synaptic blockers on the surface of the cerebellum, instead of trying to inject them deep into the tissue, has the advantage of preserving the integrity of the tissue but the disadvantage of relatively slow diffusion.

**Figure 3:**
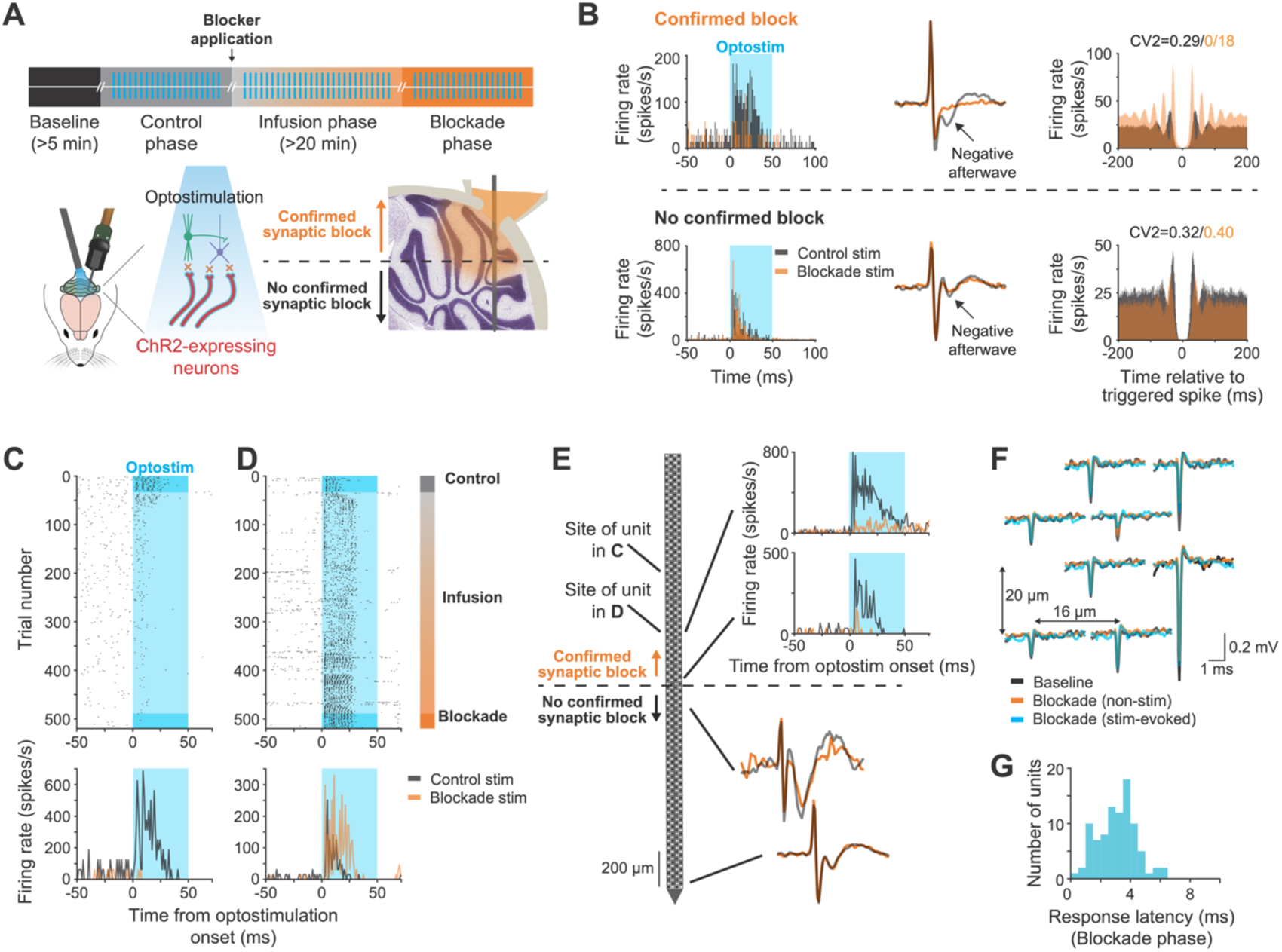
Strategy for ground-truth identification of cell type. A. Schematic showing the sequential phases in an experiment designed to test for optogenetic activation in the presence of synaptic blockers. B. Examples of the results used to verify the region of synaptic blockade. Examples above versus below the horizontal dashed line were taken as evidence for versus against blockade at that site. From left to right, we assayed the effect of blockade on the response to optogenetic stimulation, the negative afterwave of a putative mossy fiber waveform, and the discharge statistics defined by autocorrelograms and the value of CV2. C. Raster and peri-stimulus time histogram for a neuron that lost its response to optogenetic stimulation with synaptic blockade. Trial numbers on the y-axis align with the cartoon showing the periods in the experiment to the right of D. Black versus orange histograms show responses before versus during synaptic blockade. Blue shading indicates the time of photostimulation. D. Same as C except for a neuron that retained its response to optogenetic stimulation during synaptic blockade. E. Example of how we determined whether the recordings in C and D were within the region of synaptic blockade. The cartoon schematizes a Neuropixels probe, the top histograms on the right show sites that were within the region of blockade because they lost their responses to optogenetic stimulation, and the lower waveforms show mossy fibers that were below the region of blockade because they retained their negative afterwaves. F. Spatial footprint of the neuron in D. Black, orange, and blue traces show the similarity of the waveforms recorded during the baseline period, during synaptic blockade without optogenetic stimulation, and during synaptic blockade with optogenetic stimulation. G. Distribution of neural response latencies to optogenetic stimulation of directly-activated neurons in presence of synaptic blockade.

We accepted neurons as activated directly by photostimulation only if we had strong evidence that they were within the locus of successful synaptic blockade and they continued to have reliable, short-latency responses to light. To determine whether synaptic blockade was effective at a given recording depth, we evaluated recordings at or below that depth and looked for any of the indications in the top row of Figure 3B:

1. Loss during the *blockade phase* of responses to optogenetic stimulation present in the *control phase* (Figure 3B, top row, left). Neurons were excluded from the ground-truth library if they retained their response in the *blockade phase* but were outside the region of synaptic blockade (Figure 3B, bottom row, left).
2. Putative mossy fibers with loss of negative afterwaves in the *blockade phase* (Figure 3B, top row, middle). Here, we rest partly on prior evidence that the negative afterwave is a post-synaptic response of granule cells^50,51^ and that analogous negative afterwaves have been shown to correspond to post-synaptic responses in other brain regions^52^. Also, our finding of the effect of synaptic blockade on the negative afterwave in recordings from single putative mossy fibers provides the strongest evidence to date that the negative afterwave represents post-synaptic depolarization.
3. Substantial changes in a neuron’s autocorrelogram or “coefficient of variation 2” (CV2), usually due to an increase in regularity caused by a shift from synaptically-and intrinsically-driven spiking to purely intrinsically-generated spiking^31^ (Figure 3B, top row, right).

To illustrate our strategy, we provide detailed examples from one experiment in a transgenic mouse line that expresses ChR2 in mossy fibers^37,53^ (Thy1-ChR2 line 18). Here, some neurons lost their responses to optogenetic activation with synaptic blockade (Figure 3C), while others recorded nearby retained their responses (Figure 3D). We evaluated the effect of synaptic blockade along the electrode penetration that yielded these two units (Figure 3E, region above the dashed line on the Neuropixels schematic) to confirm that the neural responses in Figure 3D were activated directly by optogenetic stimulation. At sites near and deeper than the neuron in D, neurons lost their responses with synaptic blockade, indicating that they were within the region of successful blockade. Still deeper in the penetration, we recorded two putative mossy fiber waveforms that retained their negative afterwave with synaptic blockade, indicating that they were outside the region of successful blockade. Finally, the extracellular waveforms of the activated neuron were constant across the entire experiment (Figure 3F), indicating that we had a stable recording. Therefore, we concluded that the neuron in Figure 3D was an optogenetically-activated neuron, namely a mossy fiber.

Ground-truth neurons generally responded to optogenetic stimulation with latencies shorter than 5 ms (Figure 3G). However, we recommend against using latency as the sole criterion to accept or reject neurons as optogenetically-activated^21^. Latency is strongly dependent on illumination intensity and the density of opsin expression. We frequently observed short-latency responses to optogenetic stimulation that disappeared with synaptic blockade, for example the neuron in Figure 3C where the latency was less than 3 ms. Also, the neuron in the top row of Figure 3B responded to optogenetic stimulation with a latency shorter than 5 ms, but lost its response with synaptic blockade, indicating that it was driven by synaptic activation rather than direct optogenetic stimulation.

### Strategy to mitigate off-target expression in transgenic mouse lines

To varying degrees, off-target expression is a common feature of transgenic mouse lines. Often, there is no ‘clean’ line available where expression is limited to a given cell-type of interest. Our strategy to obtain ground-truth cell-type identification despite off-target expression was to (i) characterize the anatomical specificity of opsin expression for all mouse lines under study and (ii) combine identification of the recording layer based on Phyllum with optogenetic activation in the confirmed presence of synaptic blockers to establish cell-type unambiguously.

The problem of off-target expression was most pronounced in the GlyT2-Cre line used previously to image activity in Golgi cells^35^. The GlyT2-Cre line has substantial off-target expression in molecular layer interneurons and, very occasionally, Purkinje cells (Figure 4A, Supplementary Figure 2): the relative density of molecular layer interneurons was higher than that of Golgi cells (Figure 4B, 79 vs. 20%). Accordingly, we recorded neurons directly responsive to optogenetic stimulation in both the granule cell layer and the molecular layer (Figure 4C). We used Phyllum to identify the recording layers and labeled units in the granule cell layer that were directly activated by optogenetic stimulation as Golgi cells (Figure 4C, green PSTH). We labeled activated units in the molecular layer as molecular layer interneurons (Figure 4C, magenta PSTH).

**Figure 4:**
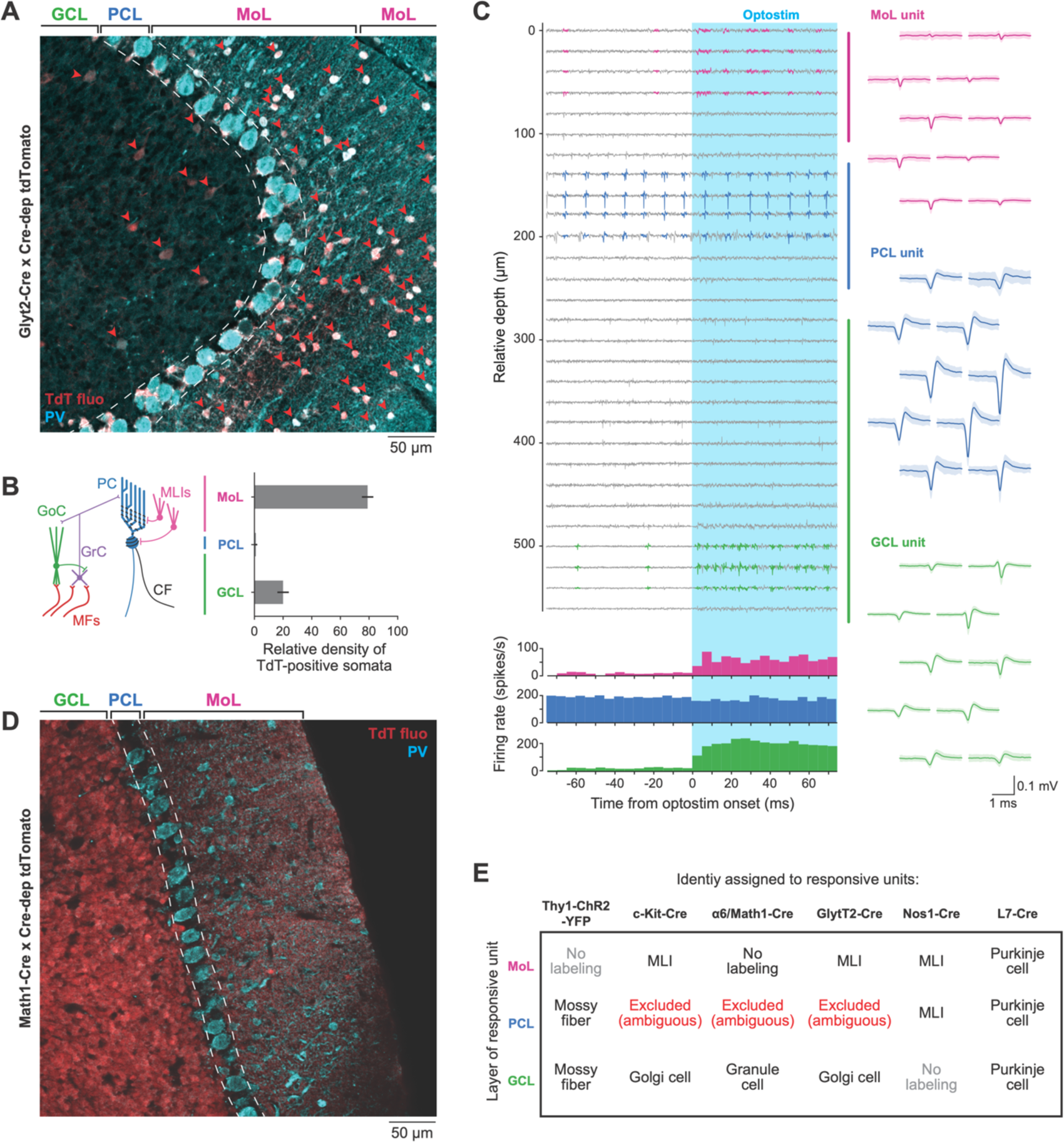
Analysis and mitigation of off-target expression in mouse optogenetic lines. A. Double stained section of cerebellum in the GlyT2-Cre line showing expression in both Golgi cells in the granule cell layer and molecular layer interneurons. Red arrows point to cells that express Td-Tomato. Blue cells express parvalbumin (PV). MoL, molecular layer; PCL, Purkinje cell layer; GCL, granule cell layer. B. Cartoon of cerebellar circuit and histogram showing density of TdT-positive somata in each of the three layers in a GlyT2-Cre mouse: GoC, Golgi cell; GrC, granule cell; PC, Purkinje cell; MLI, molecular layer interneuron; CF, climbing fiber; MF, mossy fiber. C. Representative recordings from a Neuropixels probe using optogenetics to activate neurons that express opsins in the GlyT2 line. Magenta, blue, and green waveforms on the right show the spatial footprint of neurons in the MoL, PCL, and GCL. Histograms below the voltage traces show that both the MoL and GCL layer neurons were activated by optogenetic stimulation at the time indicated by the blue shading. D. Same as A, but for the Math1-Cre line. E. Table outlines how we used layer information to disambiguate cell types despite some off-target expression in certain Cre-lines.

Other mouse lines also showed some off-target expression. For example, the Math1-Cre line used to label granule cells was generally specific (Figure 4D) but exhibited rare labeling of Purkinje cells. The c-kit-Cre line also labeled a small number of Golgi cells^34^, and the Nos1-Cre line exhibited occasional labeling of non-neuronal cells in addition to molecular layer interneurons (Supplementary Figure 2). By contrast, other lines we used were cleaner, such as the Thy1-ChR2-YFP line 18 and Pcp2-Cre lines used to label mossy fibers and Purkinje cells, respectively (Supplementary Figure 2). Crucially, we did not observe multiple labeled cell-types within a single cerebellar layer in any of our lines. Thus, the combination of an identified layer with direct optogenetic activation (Figure 4E) allowed us to disambiguate cell type for all experiments.

### The ground-truth library

Across 188 Neuropixels recordings in two laboratories, we recorded a total of 3652 neurons that survived the spike-sorting and curation pipeline (Figure 5A). Of these, 562 exhibited a response to optogenetic stimulation but only 97 passed our rigorous criteria for direct rather than synaptic activation based on reliable, short-latency responses in the presence of synaptic blockade. We added the simple spikes and complex spikes of 62 Purkinje cells identified by a complex-spike triggered pause in simple spikes. We removed 6 units recorded in Cre-lines with off-target expression where the layer of the recording was ambiguous, and 13 units that, on final closer inspection, did not have sufficiently long baseline periods due to intervals that violated our missed/extra spikes criteria (Figure 2E, F). The resulting ground-truth library contained 202 units: 69 Purkinje cell simple spikes, 58 Purkinje cell complex spikes recorded at the same time as the simple spikes, 27 molecular layer interneurons, 18 Golgi cells, and 30 mossy fibers. For comparison with previous reports^20,54,55^, Supplementary Figure 4 provides the electrophysiological signatures of different cell types in our ground-truth library of cell types using a range of metrics.

**Figure 5:**
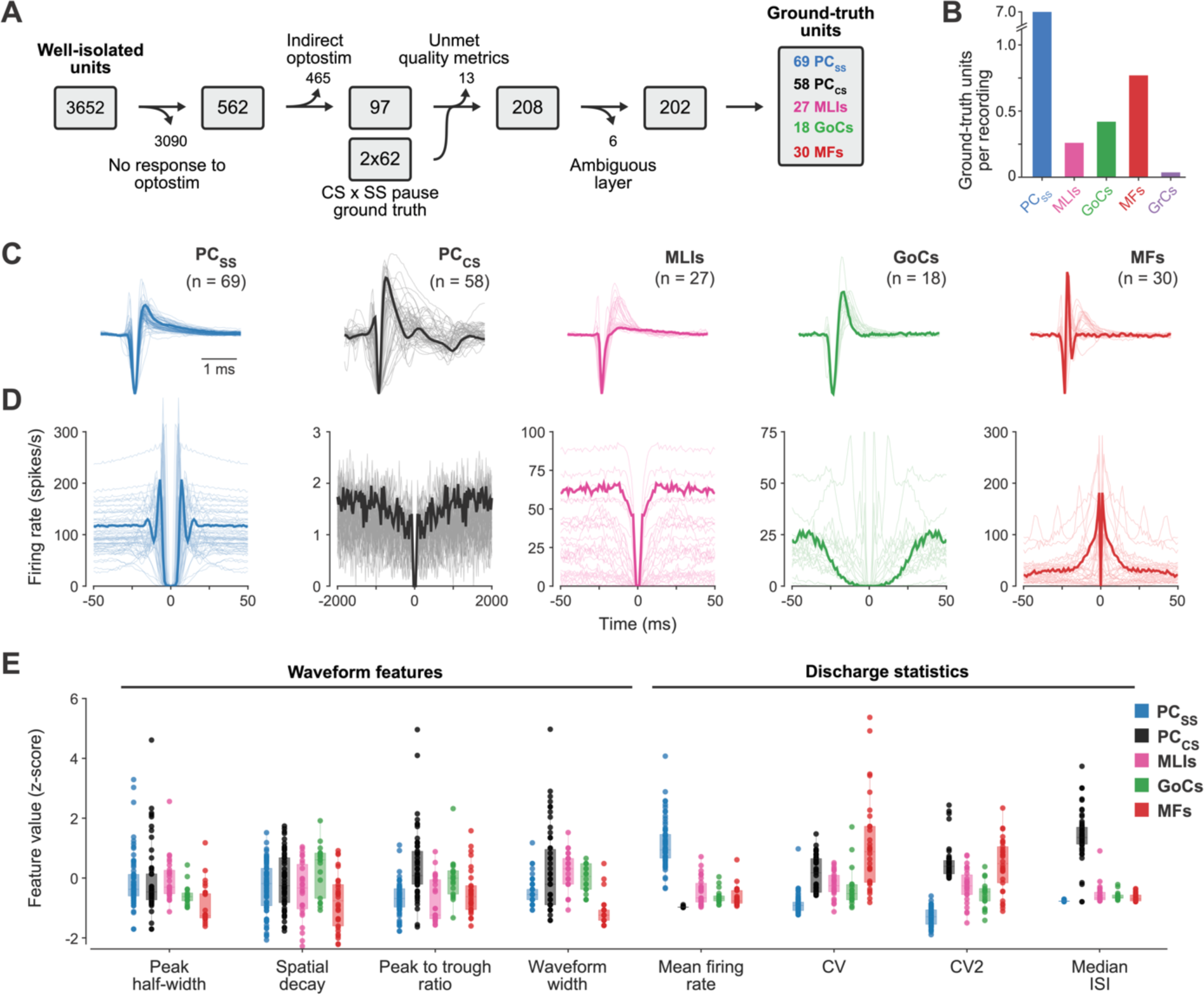
Selection criteria and properties of the ground-truth library of cerebellar cell types. A. Curation criteria used to decide which neurons to include in the ground truth library, including the numbers that were retained or deleted at each stage of the curation. B. Histogram showing the number of ground-truth units of each cell type normalized for the number of recordings: MLIs, molecular layer interneurons; GoCs, Golgi cells; MFs, mossy fibers; GrCs, granule cells. C. Superimposed waveforms for each cell type in the ground truth library. Abbreviations as in B, plus: PCSS, Purkinje cell simple spikes; PCCS, Purkinje cell complex spikes. The bold trace indicates the neuron that has an example 3D-ACG in Supplementary Figure 5. Waveforms are normalized and flipped to ensure the largest peak is negative (see ***<u>Methods</u>***). D. Same as C but showing autocorrelograms of ground-truth neurons. Note that the spike counts in the autocorrelograms have been divided by the width of the bin so that the y-axis is in spikes/s. E. Failure of traditional measurements of waveform or discharge statistics to differentiate cell types. Each symbol shows Z-scored values of different features from a single neuron; different colors indicate different cell types, per the key in the upper right. Z-scores were computed separately for each feature but across cell types within each feature. Abbreviations as in B.

In an attempt to obtain ground-truth recordings from granule cells, we made 82 recordings with Neuropixels probes in mice with the Math1-Cre or BACα6Cre-C lines either crossed to Cre-dependent ChR2 (Ai32) or injected with AAV to confer ChR2 expression (see ***Methods***). We did record multiple unit activity that was responsive to photostimulation in the region of confirmed synaptic blockade, but almost all putative single units found by KiloSort failed one or more of our criteria for good isolation (Figure 2). After careful curation, we retained zero likely granule cells from 32 recordings in the Hull lab and at most 3 from 50 recordings in the Häusser lab. The yield of fewer than 0.04 granule cells per recording was significantly lower than for the other cell types (Figure 5B). We conclude that it is challenging to record from granule cells using the current generation of Neuropixels probes; our sample is far too small to include them in the classifier we will develop next. A combination of factors may contribute to the inability to record regularly from granule cells: their comparatively small size^56,57^, the likelihood that they generate a spatially-restricted closed-field extracellular potential, and the low electrode impedance^58^ of Neuropixels^16^ (150 kOhms).

Armed with a ground truth dataset, the next challenge was to develop an accurate classification method based on consistent differences in electrophysiological features across cell types^59^. To maximize the success of classification, we strove to use both waveform^20,60,61^ and discharge statistics^62–64^ as features for cell-type classification.

> *<u>Waveform</u>*: We anticipated that the different cellular properties and morphology of different cell types would lead to different waveforms^60,61,65,66^. Patch clamp recordings *in vitro* confirmed that biophysical differences across neuron classes are manifest as consistent variations in the shape of the waveform (Supplementary Figure 3). Yet, as shown by the comparison of Figure 5B with Supplementary Figure 3, waveforms are much more variable in extracellular recordings *in vivo* than *in vitro*, and waveforms alone do not cleanly distinguish cell type.

> *<u>Discharge statistics</u>*: It is common for different cell types to have different discharge statistics throughout the brain^67,68^ and the same is true in the cerebellum of anesthetized animals^62–64^. In awake animals, discharge statistics are likely to vary across cerebellar regions and to depend on the specific behavior or sensory input^20,69^ . Therefore, a robust classification strategy should harness additional information that normalizes for the factors that contribute to variation in awake animals.

### Cell-type identification from a semi-supervised deep-learning classifier

Our deep-learning classifier strategy takes advantage of the rich information contained in the diverse waveforms and firing statistics in the ground-truth library (Figure 5C, D), along with the layer information that also provides information about cell type. Rather than using a potentially biased set of investigator-chosen measurements from waveform and firing statistics, we chose to use raw data because they (i) contain richer information, (ii) provide unbiased inputs for cell-type identification, and (iii) are likely to generalize across regions, tasks, and species. Further, Figure 5E and Supplementary Figure 4 reveal that it is difficult to guess which specific measures of waveform and firing statistics would be most informative to successfully distinguish cell types in awake animals.

We represent spike waveforms as the full time-course of the average, drift-and shift-corrected waveform on the channel with the largest signal. We represent firing statistics as autocorrelograms (ACGs) that assess the firing rate of a neuron as a function of time relative to each spike. Because the traditional two-dimensional-ACG (2D-ACG) is subject to artifacts when neural firing rate varies across a recording session or in relation to behavior, we developed “three-dimensional autocorrelograms” (3D-ACGs) that normalize for firing rate (Figure 6A, see ***Methods***). Supplementary Figure 5 shows example 3D-ACGs for each ground-truth cell type. We developed a strategy to avoid the potential issue of overfitting that is inherent in a deep-learning classifier given the high dimensionality of the waveforms and 3D-ACGs and the relatively small number of training examples in the ground-truth library. To address the mismatch of training data relative to input dimensionality, we used an unsupervised dimension-reduction technique^70^ that took advantage of 3090 unlabeled single units recorded with Neuropixels probes during the optogenetics experiments. We trained two variational autoencoders (Figure 6B), one each for waveforms and 3D-ACGs, to reduce the input dimensionality to 10 for both features (see ***Methods***), thereby minimizing the number of parameters in the ultimate classifier that needed to be trained *de novo*.

**Figure 6:**
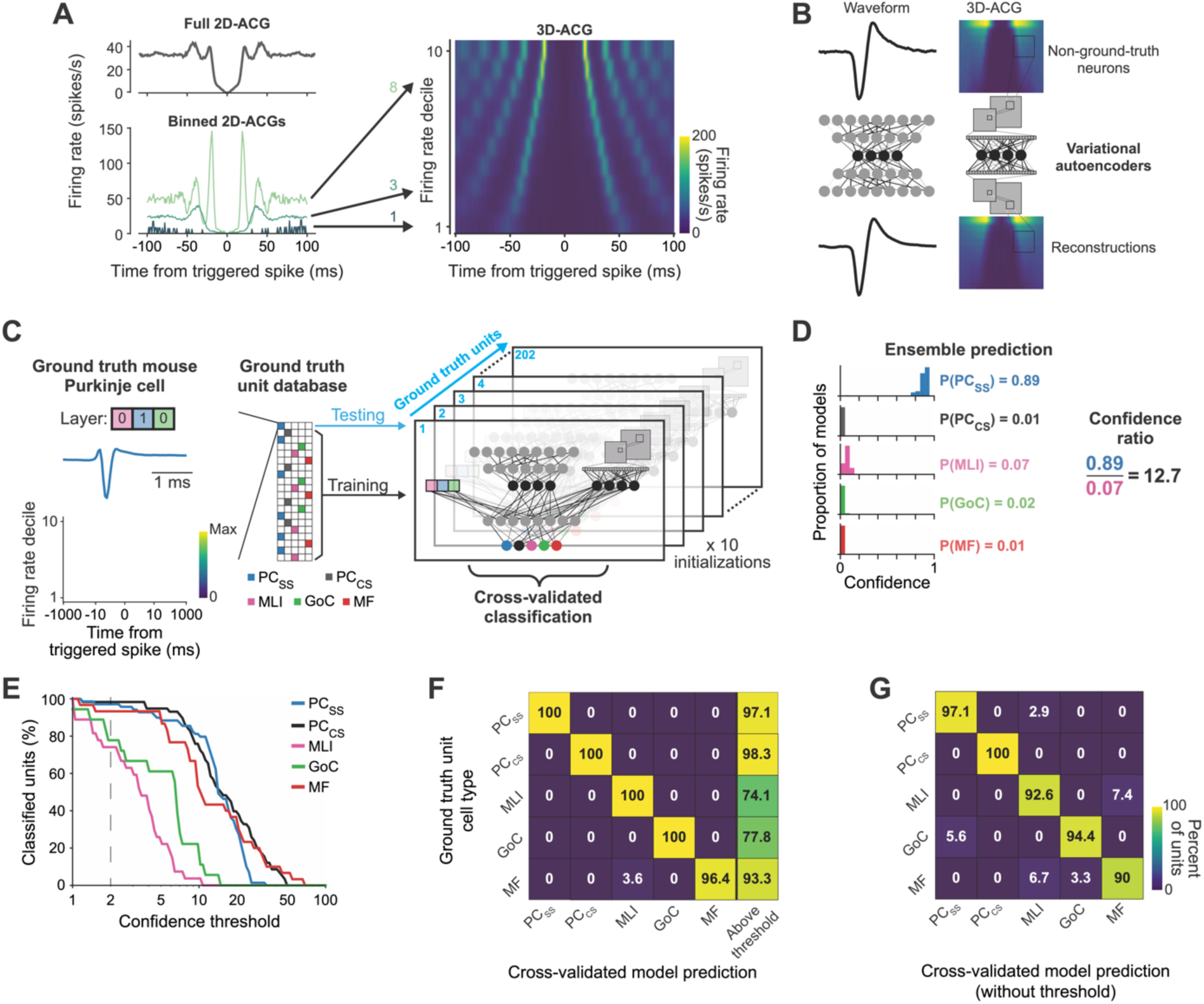
Performance of a deep-learning classifier on cell type identification for the ground-truth library. A. Method for normalizing effects of mean firing rate on firing statistics through three-dimensional autocorrelograms (3D-ACGs). Left graphs show the consensus ACG for an example neuron without regard for firing rate on top and 3 ACGs for different mean firing rates on the bottom. The heatmap on the right plots 10 rows that show 2D-ACGs as heatmaps for 10 different deciles of mean firing rate. Arrows indicate the row in the 3D-ACG for each 2D-ACG. B. Schematic of autoencoders used in unsupervised learning to reduce the dimensionality of the waveform and 3D-ACG inputs to the classifier. C. Classifier architecture. Note that we ran the classifier with 10 different initializations for each of the 202 ground-truth units, symbolized by the 202 pages in the classifier. D. Histograms showing the predictions of the classifier on 10 repetitions of training starting with different initial conditions to develop an estimate of confidence from the means of the probabilities assigned to each cell type. E. Percentage of units classified as a function of the ratio we chose as a threshold for confidence in the assignment of cell type. Different colors show data for different ground-truth cell types. F. Confusion matrix showing the agreement between the predictions of the classifier on a single left-out testing unit and the ground-truth cell type of that testing unit. The numbers in each cell indicate the percentage of ground-truth cell types on the y-axis for each prediction of the classifier on the x-axis, where confidence was required to be greater than 2. The rightmost column shows the percentage of ground-truth neurons that received a confidence greater than 2. G. Same as F, but for neurons in the ground-truth library regardless of confidence, i.e. confidence threshold = 0.

Our classifier (Figure 6C) consists of: (i) a multi-headed, normalized input layer that accepts the 10-dimensional representations of the waveform and 3D-ACG produced by the variational autoencoders, along with a “one-hot” 3-bit binary code of the unit’s cerebellar layer; (ii) a hidden layer that processes the 3 normalized inputs, and (iii) an output layer with one output unit for each of the 5 cell types. The value of the output units sums to 1 so that the output of the classifier is the probability that a given set of inputs are from each of the 5 cell types. We trained the weights in the classifier on the data in the ground-truth library using gradient descent with a leave-one-out cross-validation strategy.

We evaluated not only the accuracy, but also the “confidence” of the output from the classifier. For each leave-one-out sample (n = 202 ground-truth units), we trained an ensemble of 10 models with random initial conditions. We then averaged the classifier-predicted probability for each cell type across model instantiations. For the example illustrated in Figure 6D, the distributions of cell-type probability reveal repeated predictions that the held-out unit was a Purkinje cell simple spike. The average probability assigned to the simple spike was 0.89 while the average probability assigned to each of the other cell types was less than 0.1. However, that need not have been the case: if the data for a given unit were compatible with more than one cell type, then the classifier might classify the unit as highly-probable to be cell type #1 in one model instance and highly-probable to be cell type #2 in another instance: the average probabilities across 10 runs of the classifier might be similar and therefore closer to 0.5 for these two cell types, indicative of low classifier confidence.

We quantified the classifier confidence for each neuron with the “confidence ratio”, computed as the ratio of the mean probability of the most-likely cell type to the mean probability of the second-most-likely cell type. As expected, the percentage of ground-truth units that could be classified decreased as a function of the value of the confidence ratio we chose as the confidence threshold (Figure 6E). Classifier confidence in general was higher for Purkinje cell simple spikes, Purkinje cell complex spikes, and mossy fibers compared to Golgi cells or molecular layer interneurons. Higher confidence thresholds increase the likelihood that cell-type classification is correct, but also decrease the number of units that get classified. We chose a confidence threshold of 2 in the remainder of our analysis because it allowed the majority of neurons to be classified while providing excellent cross-validated classification performance.

The classifier showed impressive accuracy when applied to the units in the ground-truth library. For each held-out neuron that exceeded the chosen confidence threshold, we assigned it the cell type that had the highest probability, averaged across the 10 classifier runs. The classifier assigned cell types to 78% of ground-truth molecular layer interneurons and 74% of ground-truth Golgi cells at a confidence threshold of 2 (rightmost column of Figure 6F), almost all correctly as demonstrated by the values of 100% along the diagonal of the confusion matrix (Figure 6F). The classifier exceeded the confidence threshold for more than 90% of mossy fibers, Purkinje cell simple spikes, and complex spikes and again it classified nearly all such units correctly. The accuracy of the classifier degraded without a confidence threshold, but still performed quite well: it exceeded 90% accuracy on all cell types (Figure 6G). The fact that the classifier was more accurate when we required higher confidence means that 1) the classifier has a good internal model of true neuron classes and 2) the choice to set a confidence threshold improves the performance of the classifier. We note that there is some confusion between neurons in different layers despite the use of layer as an input because their waveforms and/or autocorrelograms look ambiguous to a classifier that makes a statistical prediction based on multiple inputs.The classifier performed less well without layer information, mainly because of greater conflation of Golgi cells and molecular layer interneurons (Supplementary Figure 6). Molecular layer interneurons exhibited two distinctly different waveforms with either a small or a large repolarization phase. The latter were nearly indistinguishable from the waveforms of Golgi cells. The two types of waveforms in molecular layer interneurons do not map onto the known subtypes of molecular layer interneurons^71^.

### Classifier validation of expert-labeled datasets

We next evaluated how well the ground-truth classifier (Figure 7A) generalized by attempting to predict the cell type for neurons in a sample of expert-classified, non-ground-truth recordings from mice (Medina lab) and from the monkey’s floccular complex (Lisberger lab).

**Figure 7:**
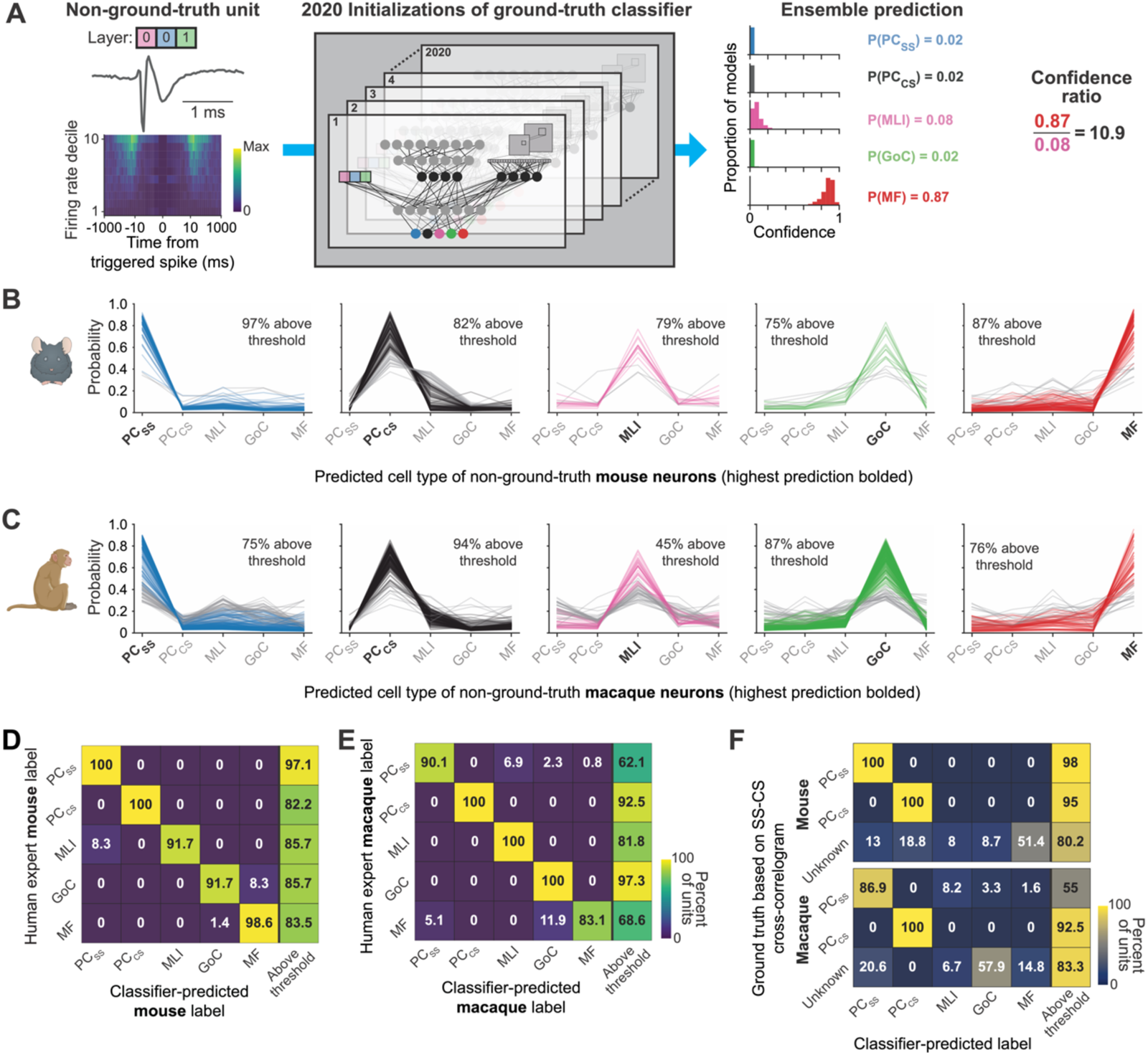
Ground-truth classifier performance on expert-classified datasets from mice and monkeys. A. Schematic of the ground-truth classifier, repeated from Figure 6C, but now making predictions based on non-ground-truth data from mouse or monkey. The n = 2020 instantiations of the classifier arise from training the classifier 10 times with different initial conditions for each of 202 left-out ground-truth units: 10 x 202 = 2020. B. Probability as a function of cell type for expert-classified neurons from mice, divided according to the cell type assigned the highest probability by the classifier. From left to right, the highest-probability cell type was a Purkinje cell simple spike (PCss), Purkinje cell complex spike (PCcs), molecular layer interneuron (MLI), Golgi cell (GoC), and mossy fiber (MF). Colored versus gray traces represent neurons that exceeded versus failed the confidence threshold of 2. Probability was averaged across runs with 2020 different forms of the classifier (see Methods). C. Same as B, but for expert-classified neurons from monkey floccular complex of the cerebellum. D. Correspondence matrix showing the agreement between the predictions of the classifier on the x-axis and the expert-labeled cell type from unclassified recordings in mice. The numbers in each cell indicate the percentage of expert-classified cell types on the y-axis as a function of the predictions of the classifier on the x-axis. The rightmost column shows the percentage of expert-classified neurons that received a confidence greater than 2 from the classifier. E. Same as D, for expert classified neurons from monkey floccular complex. F. Confusion matrices showing good agreement between the output from the classifier and the ground-truth identification in mice and monkeys of Purkinje cell simple spikes and complex spikes from the presence of a complex-spike-triggered pause in simple spike firing.

Confidence is a particularly important metric for non-ground-truth data. We took advantage of the large number of differentially trained and instantiated models from the ground-truth cross-validation analysis to improve our computation of confidence in the expert-classified datasets. The ground-truth cross-validation of the ground-truth library resulted in 2020 versions of our classifier (202 ground-truth units with 10 instantiations per cross-validation). For each unit in the expert-classified datasets, we averaged the cell-type probabilities predicted by the 2020 instantiations of the classifier and created plots of the probability assigned by the classifier as a function of cell type (Figure 7B, C). Units appear in exactly one of 5 different plots, chosen according to the cell type assigned by the classifier as the highest probability, not according to the expert-assigned cell type. For example, the leftmost graph reports probability versus cell type for all units that were classified as most probable to be simple spikes of Purkinje cells, colored according to whether the confidence ratio was below or above 2 (gray versus colored lines). The collection of confidence plots in Figures 7B and C underscores the points of confusion for the classifier, perhaps due to subtle differences in the properties of probes, the behaviors performed by the animals, or the cerebellar sites of recording. For both the mouse data and the monkey data, classifier confidence was greater than 2 for the majority of units, except that only 45% of the for units classified by the experts as molecular layer interneurons in the monkey data were classified “correctly”. The gray curves in Figure 7C suggest that the classifier often conflated molecular layer interneurons and simple spikes in the monkey data, resulting in lower confidence for these neurons.

The ground-truth classifier agreed with the human experts about the cell types in mice and monkeys of almost all units that were above confidence threshold, as demonstrated by the large percentages along the diagonal in the correspondence matrices of Figure 7D and E. Further, the ground-truth classifier was quite confident about the cell-types in the non-ground-truth data, as illustrated by the high percentage of units above a confidence threshold of 2 in the rightmost columns of Figure 7D and E. Here, it is important to explain a subtle difference in the numbers in Figures 7B and C versus Figures 7D and E. The leftmost graph of Figure 7C indicates that 75% of the monkey units classified as simple spikes exceeded confidence threshold. Because the units included in Figure 7C are not necessarily classified by experts as simple spikes, 75% is not inconsistent with the 37.9% of expert-classified simple spikes that were below confidence threshold in Figure 7E. The difference in the numbers results from analysis of the space across orthogonal axes.

The ground-truth classifier identified correctly, with confidence greater than 2, the mouse and monkey Purkinje cell simple spikes and complex spikes from recordings with a complex-spike-triggered pause in simple spikes (100/86.9% and 100/100% in mice/monkeys for simple and complex spikes, respectively. Figure 7F). “Unknown” units, defined as those not identified definitively as Purkinje cells, were distributed across cell-types by the classifier, as expected given that they included recordings from all cell types.

### Similar properties within cell types across species and cerebellar regions

Three additional analyses indicate that the success of the ground-truth classifier on the expert-classified data is based on true statistical similarity of the waveforms and firing statistics of each cell type across datasets. *<u>First</u>*, the percentage of units that we classified with confidence decreased similarly as a function of the confidence threshold for the two samples of expert-classified cells (Figure 8A, thick gray and black traces) and the ground-truth data set (Figure 8A, colored traces). *<u>Second</u>*, analysis of the output of the classifier’s autoencoders revealed excellent agreement between the reduced-dimension representation of expert-classified and ground-truth data (Figures 8B). Here, the neurons in the ground-truth library plot along the y-axis and the expert-classified neurons plot along the x-axis. Warmer colors in the heatmap indicate greater alignment of the 20-dimensional vectors defined by the concatenated outputs of the two auto-encoders in the classifier. *<u>Third</u>*, inspection of the waveforms (Figure 8C) and the 2D-ACGs (Figure 8D) reveals impressive similarity across the ground-truth data, the non-ground-truth mouse data, and the monkey recordings. Here, we have included only the neurons that were classified with confidence greater than 2. The only real exception to the visual impression of similarity is a few of the 2D-ACGs for the ground-truth Golgi cells. The similarity of resting discharge properties across preparations also appears in the 3D-ACGs (Supplementary Figure 4).

**Figure 8:**
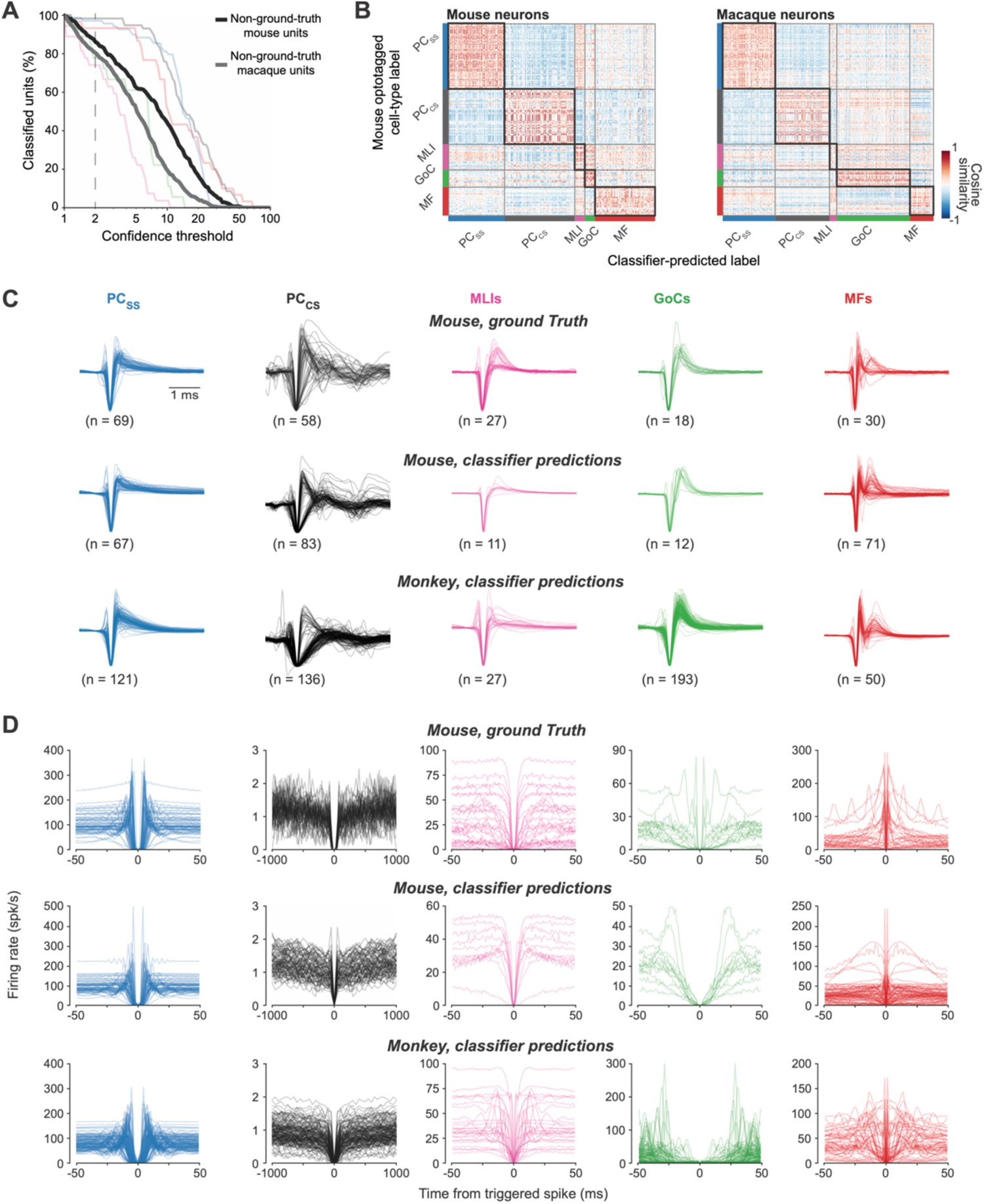
Multiple forms of evidence for the similarity of waveforms and resting discharge statistics of different cell types across the ground-truth library and the expert-labeled data from mouse and monkey. A. Comparison of percentage of classified units as a function of confidence threshold for 3 preparations. Faint colored traces show the same curves for the ground-truth library, from Figure 5E. Bold black and gray traces show results for unclassified mouse and monkey data, respectively. B. Congruence of the output from the autoencoders for identically labeled ground-truth versus expert-classified neurons across preparations. Each row corresponds to a single ground-truth identified neuron. Each column corresponds to a single classifier-identified neuron from mouse (left) or monkey (right). Colors at the intersections for each row and column indicate the cosine similarity of the concatenated outputs from the autoencoders for waveform and autocorrelograms, where redder colors indicate greater similarity. C. Waveforms of different cell-types across laboratories and species. In the first row, waveforms are divided according to ground-truth cell type in mice. In the second and third rows, cell types are divided according to classifier predictions of cell type for non-ground-truth neurons recorded in mice and monkeys. D. Same as C, except showing 2D-autocorrelograms. Note that the spike counts in the autocorrelograms have been normalized by the width of the bin so that the y-axis is in spikes/s.

The large fraction of non-ground-truth neurons that can be classified with confidence, and the agreement with the experts, is unexpected evidence that the properties of different cerebellar cell types are consistent across species and cerebellar regions. It certainly was possible, *a priori*, that a different outcome might have emerged because of genuine differences in waveform or discharge statistics across species and cerebellar regions, differences in data collection and analysis across labs, or a failure of rigor in our procedures for curating the ground-truth and expert-classified data. We anticipate that Figure 8 will serve as a useful resource for other cerebellar labs to be confident of their own rigor as they assign cell types in their own data.

### Functional dissection of cerebellar circuits enabled by cell-type identification

The ability to identify cell types from extracellular recordings allows characterization of the dynamic response properties across cell types during behavior, with the potential to reveal how a local circuit performs computation. Across our 4 labs, we were able to record and subsequently classify with confidence the principal cell types in the cerebellar circuit. We find that different neuron types often exhibit distinct temporal response dynamics during 4 different behaviors: from left to right in Figure 9A, reward conditioning in mice, eye blink conditioning in mice, locomotion in mice, and smooth pursuit eye movements in monkeys. The differences are evident in traditional peri-stimulus time histograms that summarize the average temporal profiles of each cell type (Figure 9B). Further, heatmaps with the responses of all recorded neurons of each type demonstrate considerable diversity of dynamics within each cell-type population (Figure 9C). Here, we enabled direct comparison across cell types with different absolute modulations by normalizing the response of each neuron to the standard deviation of pre-trial baseline firing. We included single-session recordings in the 3 mouse preparations and a larger pseudo-population recorded across many sessions in monkeys^72–74^. The normalization reveals that many neurons responded strongly during the behaviors, but there also were many neurons that responded weakly or not at all.

**Figure 9.**
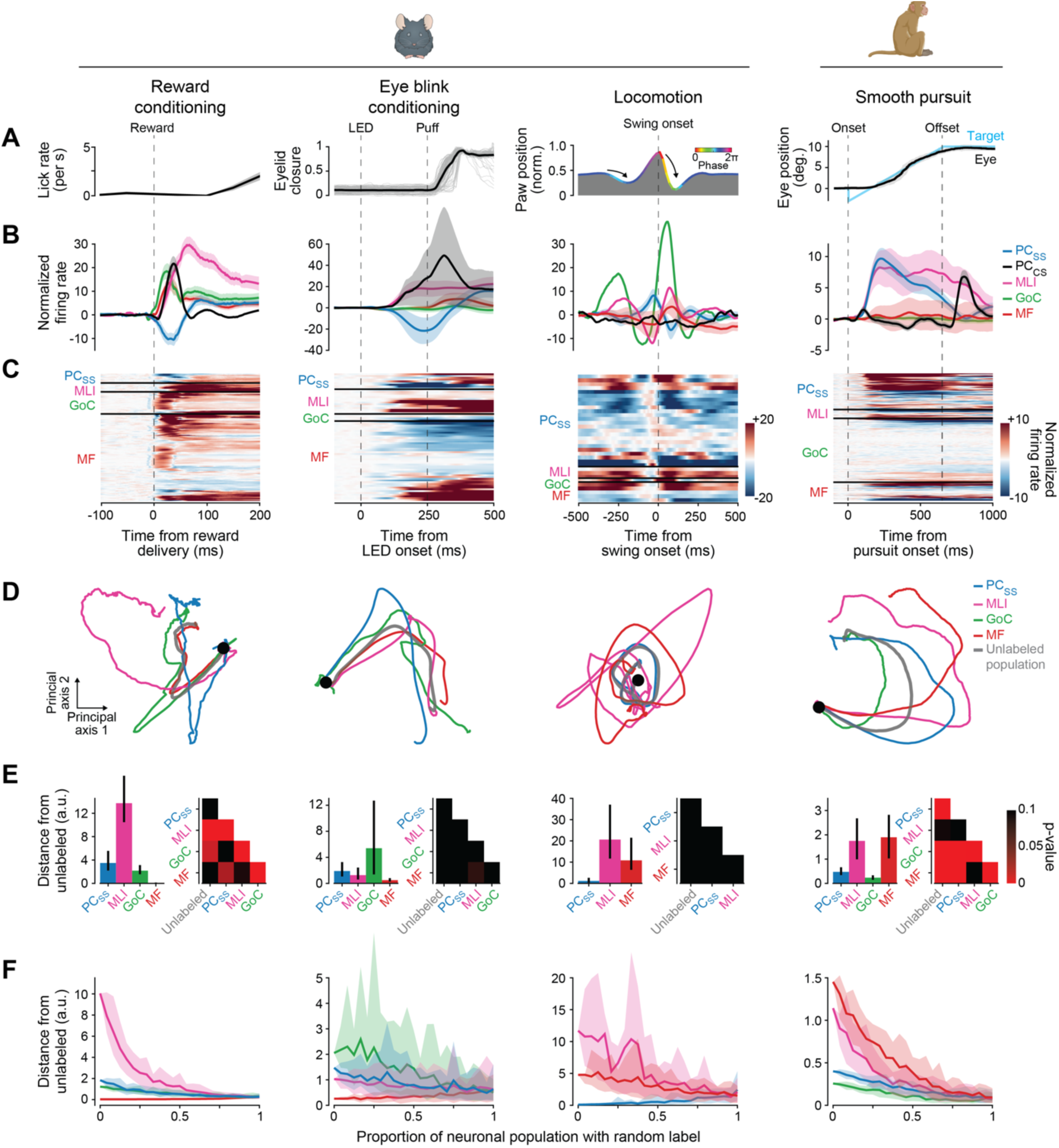
Dynamic trajectories of population responses for individual cell types. A. Temporal evolution of behavioral tasks in the 4 labs. From left to right: licks per second when a reward is cued by a light; eyelid closure when eye blinks have been conditioned by an LED as conditioned stimulus and an air puff as unconditioned stimulus; paw position aligned on the onset of swing phase during locomotion; eye position during pursuit of a step-ramp target motion. B. Average firing rate of different cell types. Different colors indicate normalized responses for the different cell types. Error bands show mean ± sem. C. Heatmaps where each line shows the normalized firing rate of one neuron, divided according to cell type, as a function of time during the four behaviors. We normalized the firing of each neuron so that the standard deviation of rate in the baseline period was 1. D. Two dimensional trajectories of population dynamics for each full population of neurons and for the individual cell types. The trajectories start at the black, filled circle, the gray trajectory shows the full, unlabeled population, and the colored trajectories show results for different cell types. E. Statistical analysis of the differences among the dynamic population trajectories of different cell types. The lefthand histograms show the distance from each cell type’s trajectories to the trajectory derived without cell-type labels. The righthand half-matrices summarize the p-values for comparison of all trajectories with each other. Black squares are not statistically significant. F. Effect of re-labeling different fractions of each cell-type population randomly on the distance between the dynamic population trajectory for each cell type and the trajectory for the unlabeled, full population.

We directly visualized the neural dynamics across the complete population of neurons recorded in Figure 9C by performing dimensionality reduction on the neural state-space using principal component analysis^72,73,75–77^. To ask whether each cell type displayed low-dimensional dynamics consistent with the full recorded population, we optimally aligned the principal components computed separately for each cell type population with those discovered across the complete population (see ***Methods***). The resultant neural trajectories for reward conditioning and smooth pursuit eye movements (Figure 9D) showed statistically different (Figure 9E) dynamic trajectories from each other and from the entire population. For eye blink conditioning and locomotion, the population dynamics failed to reach statistical significance. In all 4 systems, the differences in population dynamics between cell types collapse gracefully as a function of the fraction of cells that are labeled randomly, rather than by the classifier (Figure 9F). The graceful collapse validates that our classification of cell types at least stratifies neurons according to functional properties that can be expected to relate to cell type. Together, the analyses in Figure 9 argue strongly that cell-type identification yields biological insight about the dynamics that emerge as outputs from a given brain area and the computations performed within an area. The insight is critical for understanding how neural circuits work on the basis of interactions among different cell types.

Our analysis highlights some cautions. <u>First</u>, the “cell-type agnostic” population is biased strongly by the availability of neurons of a given type (Figure 9E). If a given class of neurons is easier to record, it will bias the ‘population response’. If the population is biased heavily towards input fibers, as in 2 of our datasets, then the cell-type specific trajectories will be essential to distinguish these dominant dynamics from those of the output neurons of an area. <u>Second</u>, failure of the trajectories of different cell types to diverge doesn’t necessarily imply that all cell types have the same dynamics. Rather, it could result from the diversity of dynamics within a cell type (owing to true heterogeneity or limited sample sizes), or the existence of 2 or more subgroups within each cell type. Non-significant trajectory differences may emerge because within group diversity is as great as between group diversity. These effects likely explain the similarity of neural trajectories within the mouse eye blink conditioning dataset.

## Discussion

Identification of cell type from *in vivo* extracellular recordings is a fundamental issue in systems neuroscience^5,6,20,55,60,62,63,68,78–81^. Our approach delivers a highly-reliable ground-truth library of the electrophysiological properties of cerebellar cell types in awake mice based on identification through optogenetic stimulation in the presence of synaptic blockers. The ground-truth library consists of the waveform of the electrical recording, the statistics of the spike train, and the layer of the cerebellum where we recorded each unit, information that comes from well-isolated neural recordings using high-density multi-contact probes. Our semi-supervised deep-learning classifier performs well in identifying the cell types in the ground-truth library, while also reporting its confidence in each identification. The internal representations in the classifier reveal similar statistics in the ground-truth library and in independent datasets of expert-classified recordings from the mouse and monkey cerebellum. The cell types predicted by the classifier for the mouse and monkey data agree with the experts’ assessments. We are encouraged by the accuracy and precision of our classifier and expect that it will be possible in the future to align the cell type obtained from extracellular recordings with that obtained from other levels of analysis, including anatomical and molecular fingerprints.

Cell-type identification enables important biological insights about circuit organization, circuit dynamics, and the relationship between output dynamics and behavior. For two of the behaviors we study, different cell types showed very different population dynamics, suggesting distinct roles for different cell types in cerebellar processing and circuit computation. For the other two behaviors we study, different cell types had statistically indistinguishable population dynamics, highlighting the variable neural response profiles within each cell class that underlie these behaviors. Cell-type identification opens avenues to use multiple different strategies for biological insights that will emerge from deep, systematic analysis in future papers. For example, the combination of cell-type identification and simultaneous recordings opens up the possibility of direct measurements of connectivity strengths and circuit function using cross-correlogram analysis, as well as localization of multiple sites of learning in a single circuit. Alternatively, analysis of the dynamics in different cell types could reveal population vectors related to motor behavior in output neurons and to context in specific types of interneurons.

The strategy we developed may be more useful and important than the exact classifier. Our goal at the outset of our project was to achieve cell-type identification from extracellular recordings in the cerebellar cortex across laboratories and species. We think that the strategies inherent in our classifier, and the classifier itself, can be used with confidence by any cerebellar recording lab that is curating their electrophysiological data with sufficient rigor. However, we also point out that a failure of rigorous curation will lead to noisy and unnecessarily variable inputs to the classifier and will contaminate the cell-type identifications provided as its output^61^.

Others have attempted to identify the distinct signatures of discrete populations of cerebellar cortical neurons^20,62–64,82,83^. Past attempts identified either (i) neurons of interest by qualitative agreement with spiking signatures found *in vitro*^64^ or (ii) recordings in anesthetized preparations with neurons identified anatomically via juxtacellular labeling^20,58,62,63,82,83^. Our recordings in awake, behaving mice demonstrate large variance in the discrete metrics used for summarizing spiking activity both within and across ground-truth classes (Figure 5E, Supplemental Figure 4). Thus, classification schemes reliant on a finite set of specific features probably will not generalize well to other tasks or regions^60^, or from anesthetized to behaving preparations^20^. We think our approach is more likely to generalize because it leverages the informativeness of the full waveform^60,61^ and 3D-ACG to classify cell types across a wide range of behaviors and stimulus-driven responses.

Several features of the strategy embedded in our classifier were critical to its success:

● *<u>Raw waveforms and 3D-ACGs</u>*. Raw features are an unbiased input^60^ and allow the classifier to take advantage of extensive information in waveform^60,61^ and discharge statistics. 3D-ACGs normalize for variations in firing rate and create a statistic that can be compared across cerebellar areas, experimental tasks, and species. Similarly, the choice to use single-channel waveforms allows the classifier to generalize across electrode types. We think our strategies are likely to generalize because they use raw features that can be measured readily in other brain areas.
● *<u>Mitigation of overfitting</u>*. We developed a semi-supervised^84–86^ deep-learning strategy (see ***Methods***) to train the classifier with a relatively small number of ground-truth neurons. The unsupervised training of variational autoencoders reduces the chances of overfitting^87^. The use of a large unlabeled dataset to train the autoencoders also reduces the chances of overfitting by ensuring that their architecture was designed independently from the ground-truth dataset. Successful predictions of cell type that agree with two expert-classified datasets supports the generalizability of the classifier on other data. Our choice of how to mitigate overfitting should allow our strategy to generalize to other datasets where the dimensionality of the inputs is high and the number of ground-truth neurons is comparatively small.
● *<u>Confidence</u>*. We were particularly cognizant of making our classifier trustworthy. To do so, we established confidence by training multiple models on the same data^88^ and by using a Bayesian method to calibrate confidence at the single model level^89,90^ (see ***Methods***). By requiring confidence above a given threshold^91,92^, we improved the accuracy of the model on the ground-truth data as well as for non-ground-truth recordings. The ability to choose a confidence threshold allows the user to balance whether to include all neurons even if some cell type assignments might be incorrect or to include fewer neurons with greater certainty in the cell type assignment. Reliable spike sorting and cell-type classification is challenging with multi-unit recording techniques. Thus, it is important that population dynamics collapse across cell-types when they are labeled randomly. The importance of cell-type labels implies that we successfully partitioned a large proportion of the different cell types. and that our conclusions are not undermined by any errors in cell-type identification.

The classifier was more successful when it included layer information as an input. With layer information, it classified a higher percentage of the units and, in the ground-truth data, classified them with greater accuracy. However, the use of layer as an input does not make classification trivial. Rather, it creates a platform that will become even more useful as we are able to achieve ground-truth identification of other cell-types in the cerebellum, for example of granule cells with improved recording probes. Also, because waveform and firing statistics are necessary to distinguish cells that are in the same layer, the classifier makes a statistical decision about cell type rather than relying solely on layer for cell identification^93^. Layer is defined in a specific way for the cerebellum^1^, but we think of layer information more generally as a specific example of “local electrical properties”. We imagine that there are other ways to quantify those properties, for example LFPs and current-source-density analysis^94,95^, that will work in brain areas without a laminar structure.

The biggest challenge, and our bigger goal, is deployment of the strategy outlined here in other brain areas. The use of layer information to improve classification should be relevant to other structures – cerebral cortex^96^, hippocampus^97^, superior colliculus^98^ – that have layers with measurable local electrical properties. We also think that the strategies used in our classifier enable generalization by showing how to reduce the dimensions of raw data used as inputs while mitigating the challenges of small numbers of neurons in training sets. We hope that application of our strategy in other brain areas will enable cell-type identification from extracellular recordings, a key element in our collective long-term goal of understanding how neural circuits work and how they generate behavior.

### Limitations of the study

One class of limitations is related to our procedures for data collection and curation. While we were scrupulous about spike sorting and criteria for data inclusion, we relied on indirect measures and logic to determine whether a recording was within the region of synaptic blockade. Any errors might have allowed inclusion of certain mis-identified cell types due to off-target opsin expression. We assessed off-target expression thoroughly, revealing, for example, extensive off-target expression in the GlyT2 line^35,99,100^. Less obvious off-target expression could have escaped our histological analysis and introduced a small fraction of incorrectly-identified cells, for instance in the Math1 line^101^.

A second class of limitations is related to the ground-truth classifier’s performance for labeling neurons outside of those we focused on in this study. While we trained the classifier on 5 identified cell types, other neuronal subtypes exist in the cerebellar cortex^28^. We may have recorded from the rare cell types in our expert-labeled datasets. Were they misclassified as one of the 5 cell types in our ground-truth library? Or were they correctly recognized as “other” cell types and relegated to the ∼15-20% of neurons that failed to reach confidence threshold? In due course, we expect to be able to augment our ground-truth library with unipolar brush cells^102^, candelabrum cells^103,104^, and other Purkinje layer interneurons^105,106^ as ever more specific Cre lines and viruses become available. We anticipate that advancements in recording probes with higher impedance and/or more dense recording sites will enable reliable recordings from granule cells. If waveforms, firing properties, and layer prove insufficient to segregate identity as other cell types are incorporated, we anticipate that additional information about synaptic and electrical connectivity can further improve the accuracy of our classifier.

A third possible limitation is related to the electrophysiological features we chose as inputs for our classifier. Because our goal was to identify cell type based on extracellular recordings with increasingly popular high-density probes, we used waveform, discharge statistics, and layer as inputs. The use of discharge statistics might be problematic in structures with little or no spontaneous discharge in some cell types, although 3D-ACGs enable assessment of discharge statistics from firing related to sensory inputs or behavior. Also, our use of physiological characteristics as inputs to the classifier limits our ability to align electrophysiological cell-type identifications with those provided by single-cell RNAseq^107–109^, juxta-cellular labeling^62,63^, or combinations of *in vivo* recording and single-cell imaging^110^. For example, extracellular electrophysiology cannot ‘mark’ recorded cells for post-hoc analysis of molecular identity using approaches such as RNAseq. However, our approach affords several advantages over recording methods that are compatible with genetic profiling but sacrifice experimental throughput, tissue accessibility, temporal resolution, and/or the number of cell-types that can be simultaneously recorded and identified. Thus, it is necessary to weigh the trade-offs of different strategies for cell-type identification according to the particular experimental question at hand.

Despite its limitations, our study demonstrates a strategy that allows different cell types to be identified robustly and reliably, even when analyzing independently collected data. Thus, the strategy should be of great value to the growing community of cerebellar researchers using high-density silicon probes. Furthermore, our strategy provides a template for principled semi-automated detection of cell type, based on assembly of a ground-truth library, that can be applied across other neural circuits in the brain.

## Methods

We conducted experiments in four laboratories and on two species, mice and macaque monkeys. All mouse procedures in the Häusser lab were approved by the local Animal Welfare and Ethical Review Board at University College London and performed under license from the UK Home Office in accordance with the Animals (Scientific Procedures) Act 1986 and in line with the European Directive 2010/63/EU on the protection of animals used for experimental purposes.

Mouse procedures in the Hull and Medina labs were approved in advance by the *Institutional Animal Care and Use Committees* at Duke University and the Baylor College of Medicine, respectively, based on the guidelines of the United States’ *National Institutes of Health*. Monkey procedures in the Lisberger lab were approved in advance by the *Institutional Animal Care and Use Committee* at Duke University. Every effort was made to minimize both the number of animals required and any possible distress they might experience.

### Mouse experimental procedures

#### Mouse lines

All transgenic mice were maintained on the C57BL/6J background. Both male and female mice were used and results were pooled.

● *Häusser:* Mice expressing Channelrhodopsin-2 (ChR2) in various cerebellar cell types were generated primarily by crossing Cre lines to a Cre-dependent ChR2-eYFP reporter line^111^ (Ai32, B6.Cg-Gt(ROSA)26Sortm32(CAG-COP4*H134R/EYFP)Hze/J), or, in a subset of experiments, by injecting Cre-dependent ChR2 virus (AAV1.CAGGS.Flex.ChR2-tdTomato [UPenn]). Cre lines were: BAC-Pcp2-IRES-Cre (B6.Cg-Tg(Pcp2-cre)3555Jdhu/J), intended to label Purkinje cells^36^; Nos1-Cre (B6.129-Nos1tm1(cre)Mgmj/J), intended to label molecular layer interneurons^112^; Glyt2-Cre (Tg(Slc6a5-cre)1Uze), intended to label Golgi cells^35^; and Math1-Cre (B6.Cg-Tg(Atoh1-cre)1Bfri/J), intended to label granule cells^101^. In addition to the transgenic crosses and viral ChR2 expression, we used the Thy1-ChR2 line 18 (B6.Cg-Tg(Thy1-COP4/EYFP)18Gfng/J) to express ChR2 in mossy fibers^37^. Recordings using each strategy were performed as follows: L7-Cre x Ai32 – 1 recording (1 mouse), Nos1-Cre x Ai32 – 40 recordings (34 mice), Nos1-Cre + AAV1.CAGGS.Flex.ChR2-tdTomato – 3 recordings (3 mice), GlyT2-Cre x Ai32 – 32 recording (31 mice), GlyT2-Cre + AAV1.CAGGS.Flex.ChR2-tdTomato – 3 recordings (3 mice), Math1-Cre x Ai32 – 47 recording (38 mice), Math1-Cre + AAV1.CAGGS.Flex.ChR2-tdTomato – 3 recordings (3 mice), and Thy1-ChR2 line 18 – 26 recordings (22 mice). The specificity of opsin expression in the cerebellum of our Cre transgenic crosses was further investigated by crossing the listed Cre lines to a Cre-dependent tdTomato reporter line, Ai9 (B6.Cg-Gt(ROSA)26Sortm9(CAG-tdTomato)Hze/J)^113^, so that we could evaluate expression specificity through cytosolic, rather than membrane-bound, fluorescence.
● *Hull:* Mice expressing ChR2 or the inhibitory opsin GtACR2 were generated by either crossing the *c-kit*^IRES-Cre^, intended to label molecular layer interneurons^34^ or BACα6Cre-C, intended to label granule cells^114^, to Ai32^111^ or a Cre-dependent ArchT-GFP reporter line, Ai40 (B6.Cg-Gt(ROSA)26Sortm40.1(CAG-aop3/EGFP)Hze/J)^115^. Alternatively, we injected the same lines with Cre-dependent viruses: (AAV1.CAGGS.Flex.ChR2-tdTomato [UPenn] and AAV1.Ef1a.Flex.GtACR2.eYFP [Duke]). In addition, we used the Thy1-ChR2 line 18 to express ChR2 in mossy fibers. Recordings using each strategy were performed as follows: *c-kit*^IRES-Cre^ + AAV1.CAGGS.Flex.ChR2-tdTomato – 8 recordings (2 mice), *c-kit*^IRES-Cre^ x Ai40 – 3 recordings (1 mouse), *c-kit*^IRES-Cre^ + AAV1.Ef1a.Flex.GtACR2.eYFP – 11 recordings (6 mice), BACα6Cre-C x Ai32 – 22 recordings (12 mice), BACα6Cre-C + AAV1.CAGGS.Flex.ChR2-tdTomato – 10 recordings (4 mice), and Thy1-Chr2 line 18 – 13 recordings (4 mice).
● *Medina:* All experiments were performed in C57BL/6J mice of at least 10 weeks of age, obtained from Jackson Laboratories.

#### Experimental preparation

To prepare mice for awake in vivo recordings, in all labs they were implanted with a headplate/headpost under isoflurane anesthesia in sterile conditions. Pre-operative and post-operative analgesia were administered, and mice were allowed to recover from surgery for at least one week before being habituated to head-fixation and prepared for recordings. Lab-specific details are as follows:

● *Häusser:* We installed a custom-made aluminum headplate with a 5 mm long and 9 mm wide oval inner opening over the cerebellum. Mice received a steroid anti-inflammatory drug at least 1 hour before surgery (Dexamethasone, 0.5 mg/kg), followed by an analgesic NSAID (Meloxicam, 5mg/kg) immediately before surgery. Anesthesia was induced and maintained with 5% and 1-2% isoflurane, respectively. The headplate was positioned over the lobule simplex of the left cerebellar hemisphere, angled at approximately 26° with respect to the transverse plane, and attached to the skull with dental cement (Super-Bond C&B, Sun-Medical). Post-operative analgesia (Carprieve, 5 mg/kg) was given for 3 days. After several days of habituation on the recording apparatus, a 1 mm-diameter craniotomy and durotomy were performed to allow access for Neuropixels probes into the lobule simplex (3 mm lateral to the midline, anterior to the interparietal-occipital fissure). Before the craniotomy, a conical nitrile rubber seal (Stock no. 749-581, RS components) was attached to the headplate with dental cement to serve as a bath chamber. The exposed brain was then covered with a humid gelatinous hemostatic sponge (Surgispon) and silicone sealant (Kwik-Cast, WPI) until the experiment was performed (1-2 h after recovery). At the beginning of the experiment, mice were head-fixed, the silicone sealant was removed, and physiological HEPES-buffered saline solution was immediately applied to keep the craniotomy hydrated.
● *Hull:* We installed a titanium headpost (HE Palmer, 32.6x19.4 mm) to the skull and a stainless-steel ground screw (F.S. Tools) over the left cerebellum, both secured with Metabond (Parkell). Mice received dexamethasone (3 mg/kg) 4-24 hours before surgery and an initial dose of ketamine/xylazine (50 mg/kg and 5 mg/kg, IP) and carprofen (5 mg/kg) 20 min before induction with isoflurane anesthesia. Isoflurane was administered at 1-2% throughout surgery to maintain appropriate breathing rates and prevent toe pinch response, which were monitored throughout the duration of the surgery. Body temperature was maintained with a heating pad (TC-111 CWE). Mice received buprenex and cefazolin (0.05 mg/kg and 50 mg/kg respectively, subq) twice daily for 48 hours after surgery and were monitored daily for 4 days. After 2+ weeks of recovery, mice received dexamethasone (3 mg/k) 4-24 hours before recordings. Craniotomies (approx. 0.5-1.5 mm) were opened over vermis or lateral cerebellum (relative to bregma: between -6.0 and -7.0mm AP, and between 1.0 and 2.8mm ML) on the first day of recording, under 1-2% isoflurane anesthesia, and were sealed between recordings using Kwik-Cast (WPI) covered by Metabond. Craniotomies could be re-opened for subsequent recordings under brief (<30 min) 1-2% isoflurane anesthesia.
● *Medina:* Preoperative analgesia was provided (5g/kg meloxicam, subq, 0.02mL 0.5% bupivacaine and 2% lidocaine, subq) and surgery was carried out under sterile conditions. Mice were anesthetized with isoflurane (5% by volume in O_2_ for induction and 1-2% by volume for maintenance; SurgiVet) and kept on a heating pad to maintain body temperature. The skull was exposed and leveled to match the stereotaxic plane before two stainless steel screws were implanted (relative to bregma: AP -0.3mm, ML ±1.4mm) to anchor the whole preparation. A custom-made stainless steel headplate was placed over the screws and the whole preparation was secured to the skull with Metabond cement (Parkell). Additionally, a craniotomy was performed (relative to bregma: AP -5.5mm) consisting of a 5x2 mm section of bone removed to expose the cerebellar vermis and the right anterior and posterior lobes. A chamber was then built with Metabond to cover the exposed bone around the craniotomy, the dura was protected with a thin layer of biocompatible silicone (Kwik-Cast, WPI) and the whole chamber sealed with silicone adhesive (Kwik-Sil, WPI). Mice were monitored until fully recovered from anesthesia and analgesia was provided during the three days following the surgical procedure.

#### Recording procedures

All labs followed the same general procedures for mouse cerebellar recordings. Mice were progressively habituated to head fixation prior to Neuropixels recordings. Recordings from the cerebellar cortex were made using Neuropixels 1.0 probes. Probes were coated with DiI, DiO, or DiD (Cat.Nos.V22885, V22886, and V22887; Thermo Fisher Scientific) by repeatedly moving a drop of dye along the probe shank using a pipette until a dye residue was visible along its entire length (∼20 passes). Probes were inserted into the brain at a speed of 1-4 µm/s while monitoring electrophysiological signals. The recording chamber surrounding the craniotomy was bathed in ACSF, with or without blockers. After each recording, the probe was removed and soaked in Tergazyme, then soaked in distilled water, and finally washed with isopropyl alcohol. After the last recording session, the brains of most mice were fixed and processed for histology to verify recording locations.

In all three laboratories, Neuropixels data were acquired using SpikeGLX (https://github.com/billkarsh/SpikeGLX). Following data acquisition, automated spike sorting was performed using Kilosort 2.0^17,116^ and manual curation was performed using Phy (https://github.com/cortex-lab/phy). Across all labs, signals were digitized at 30 kHz. Onboard filtering was turned on in some but not all cases.

#### Optogenetic stimulation and pharmacology

The same general procedures were followed for optogenetic stimulation in both the Häusser and Hull labs. This procedure consisted of four parts: (1) a baseline recording period without stimulation or drug block, (2) a period of optogenetic stimulation without drug block, (3) a period during which synaptic blockers were applied and diffused into the brain, and (4) a period of optogenetic stimulation with blockers present. The details of the procedures for ground-truth identification of cell-type varied slightly between the two labs.

● *Häusser:* Optogenetic stimulation was performed using 1 or 2 blue LEDs (470 nm, Thorlabs M470F3) and in some experiments a blue laser for surface illumination (Stradus 472, Voltran). Surface illumination was performed by coupling the laser or the LED via a patch cable (M95L01, Thorlabs) to a cannula (CFMXB05, Thorlabs) positioned in contact with the brain surface near the probe. In some experiments a second illumination source - a tapered fiber (Optogenix 0.39NA/200μm) glued directly to the head of the Neuropixels probe - was inserted into the brain. Total power at the fiber tip (surface fiber) and coupling cannula (tapered fiber) was 1-6.9 mW. Each recording session consisted of: (1) a 20 minute baseline period of spontaneous activity, (2) a set of 50 optogenetic stimuli (stimulation duration: every 10 seconds, 1 stimulation of 250 ms or a train of 5 stimulations of 50ms at 5 Hz, depending on the experiment), (3) an application of a synaptic blocker cocktail (Gabazine 0.2-0.8 mM, NBQX 0.8 mM, APV 1.6 mM, MCPG 0-1.3 mM) to the surface of the cerebellum followed by a 20 minute incubation, and (4) a second set of 50 optogenetic stimuli in the presence of synaptic blockers. We note that we did not record any neurons in the ground-truth library with the blue laser as a source of photostimulation.
● *Hull:* Neurons expressing ChR2 were activated and neurons expressing GtACR2 were inhibited using a 450 nm laser (MDL-III, OptoEngine) using a 400 micron optic patch fiber (FT400 EMT, Thorlabs) that was positioned 4-10 mm from the brain surface. Power at the brain surface was approximately 2-30 mW and was calibrated for each experiment to produce neuronal responses with minimal artifact. Laser stimulations lasted 50 or 100 ms and were delivered at 0.1 Hz throughout the recording after the 20 minute baseline period, with brief pauses to replenish ACSF or apply blockers (Gabazine 0.2-0.8 mM, NBQX 0.6-1.2 mM, AP-5 0.15-0.6 mM, MCPG 1-2.5 mM).

#### Histology

- ● *Häusser:* Mice were deeply anesthetized with ketamine/xylazine and perfused transcardially with PBS followed by 4% PFA in PBS. The brains were dissected and post-fixed overnight in 4% PFA, then embedded in 5% agarose (for electrode tract reconstruction) or sectioned at 100 μm (for immunohistochemistry). To reconstruct electrode tracts, we imaged full 3D stacks of the brains in a custom-made serial two-photon tomography microscope coupled to a microtome^117^, controlled using ScanImage (2017b, Vidrio Technologies) and BakingTray (https://github.com/SainsburyWellcomeCentre/BakingTray, extension for serial sectioning). The entire brain was acquired with the thickness of physical slices set at 40 µm and that of the optical sections at 20 µm (2 optical sections/slice) using a piezo objective scanner (PIFOC P-725, Physik Instrumente) in two channels (green channel: 500–550 nm, ET525/50; red channel: 580–630 nm, ET605/70; Chroma). Each section was imaged in 1025 x 1025 µm tiles at 512x512-pixel identification with 7% overlap using a Nikon 16x/0.8NA objective.

After slicing, samples for immunohistochemistry were blocked with 2.5% normal donkey serum / 2.5% normal goat serum / 0.5% Triton X-100/PBS for 4-6 hours at room temperature, primary antibodies for 4-6 days at 4°C, and secondary antibodies overnight at 4°C. Antibodies were diluted in blocking solution. The following antibodies were used: rat anti-mCherry (1:250, ThermoFisher M11217), Mouse anti-Parvalbumin (1:1000, Millipore MAB1572), Donkey anti-Rat-Alexa 594 (1:1000, Invitrogen), and Goat anti-Mouse-Alexa 633 (1:1000, Invitrogen). Neurotrace 435/455 (1:250, ThermoFisher N21479) was added to the secondary antibody solution. Sections were mounted and imaged on a Zeiss LSM 880 using a 20x objective in 425x425 µm tiles at 1024x1024-pixel identification.

● *Hull:* After the last day of recording, mice were deeply anesthetized with ketamine/xylezine (350 mg/kg and 35 mg/kg, IP) and perfused with PBS followed by 4% PFA in PBS. Brains were extracted and post-fixed in 4% PFA in PBS overnight, then sectioned at 100 mm using a vibratome (Pelco 102). Before sectioning, some brains were encased in a 2% agar block for stability. Slices were either stained with DAPI (DAPI, Dihydrochloride, 268298, EMD Millipore) and then mounted with mounting medium (Fluoromount-G, Southern Biotech) or were mounted with a DAPI-containing mounting medium (DAPI Fluoromount-G, Southern Biotech). Electrode tracts were visualized using a confocal microscope (Leica SP8).
● *Medina:* After perfusion with 4% PFA in PBS, brains were extracted, post-fixed in the same solution for at least 12h and then cryoprotected in 30% sucrose solution in PBS for 48h. The brains were aligned so the coronal sections would match the track angle and sectioned at 50 µm on a cryostat (Leica CM1950). Free floating sections were recovered in PBS and incubated in Hoechst solution for 3 minutes (Hoechst 33342, 2µg/mL in PBS-TritonX 0.25%, Thermo Fisher Scientific). Sections were then washed in PBS three times using fluorescence protectant medium (ProLong Diamond Antifade, Thermo Fisher Scientific). Epifluorescence was acquired at 10x magnification on an Axio Imager Z1 microscope (Zeiss), track reconstruction and measurements were made on specific microscopy analysis software (ZEN software, Zeiss).

#### Validation of ChR2 specificity

To identify the classes of cerebellar neurons that expressed optogenetic actuators, we determined the layer in which fluorescent neurons were present and whether they expressed parvalbumin (PV), which should be present in all molecular layer interneurons and Purkinje cells^118^. The location of cerebellar layers in each image were identified in the Neurotrace (fluorescent Nissl) channel. The soma locations of neurons expressing tdTomato (as a proxy for Cre expression) and PV were marked manually in grayscale images using Fiji (NIH). Neurons were deemed to express both tdTomato and PV if their somatic locations were less than 5 µm apart, and the layer of each neuron was determined by overlaying the Neurotrace laminar mask to cell locations.

### Macaque experimental procedures

Recordings in non-human primates were conducted in the *Lisberger* lab on three male rhesus monkeys (*Macaca mulatta*) weighing 10-15 kg. A portion of the primate dataset reported here have been published previously along with corresponding detailed methods^119^. Briefly, monkeys underwent several surgical procedures under isoflurane in preparation for neurophysiological recordings. In succession, we (i) affixed a head-holder to the calvarium, (ii) sutured a small coil of wire to the sclera of one eye to monitor eye position and velocity using the search coil technique^120^ and (iii) implanted a recording cylinder aimed at the floccular complex. Analgesics were provided to the monkeys after each surgery until they had recovered.

Each day, we acutely inserted either tungsten single electrodes (FHC) or, for the majority of our data, custom-designed Plexon s-Probes into the cerebellar floccular complex. Plexon s-Probes included 16 recording contacts (tungsten, 7.5 µm diameter) spaced in two columns separated by 50 µm. Adjacent rows of contacts were also separated by 50 µm. Once we had arrived in the ventral paraflocculus, we allowed the electrode to settle for a minimum of 30 minutes. We recorded continuous wideband data from all contacts at a sampling rate of 40 kHz using the Plexon Omniplex system. We used a 4^th^ order Butterworth low-pass hardware filter with a cutoff frequency of 6 kHz prior to digitization to eliminate distortion of the recorded signal by the electrical field produced by the eye coils. All recordings were performed while the monkey tracked discrete trials of smooth motion of a single black target (0.5° diameter) on a light grey background in exchange for liquid reward. All analyses of the primate neurons utilized the entire recording period and were not contingent on the animal’s behavior.

### Data processing and analysis

#### Assignment of layers with Phyllum

For recordings in the mouse, we assigned each channel of the Neuropixels probe to a layer using *Phyllum*^121^, a custom-designed plugin for the curation software *Phy*. The algorithm for layer identification in *Phyllum* starts by automatically setting ’anchor’ channels whose recorded layer can be unambiguously identified by the presence of Purkinje cell units with simple and complex spikes (Purkinje layer anchor), mossy fiber units with triphasic waveforms (granule layer anchor), or low 1-2 Hz frequency units with wide waveforms indicative of dendritic complex spikes (molecular layer anchor). Then, *Phyllum* fills in the layer of the remaining channels via an iterative procedure based on (1) proximity to the nearest Purkinje cell anchor and (2) allowed layer transitions. Every channel assigned to the Purkinje cell layer must contain at least a Purkinje cell recording within 100 μm, but the channel may also contain additional units located in the neighboring granule or molecular layers. If none of the channels between two consecutive Purkinje cell anchors contain another anchor unit, their layer is set to ‘Unknown’. On average, *Phyllum* assigns 82% of all the channels on the Neuropixels probe to a specific layer. Histological reconstruction of 21 recording tracks confirmed that for channels that are assigned a specific layer, the assignment is highly accurate: >99% for molecular layer channels, >98% for granule layer channels, and >95% for Purkinje layer channels.

#### Curation procedures

● **Mouse**: After automated sorting with Kilosort and initial manual curation with Phy, we implemented checks to ensure that the resulting clusters selected for further analysis corresponded to single units with physiological waveforms, good isolation properties, and few or no refractory period violations. Rigorous curation was especially important for our long recordings, which could have periods of good isolation intermixed with periods of drift or poor unit isolation. We divided our recordings into overlapping segments (30 seconds segments computed every 10 seconds) and computed the ‘false-positive’ and ‘false-negative’ rates in each segment. False positives were defined as spikes that fell within the refractory period of a unit (± 0.8 ms from a given spike) and termed refractory period violations (RPVs). The proportion of false-positives was estimated as the quotient between the RPV rate and the mean firing rate^46^. False negatives were defined as spikes that were not detected because they fell below the noise threshold of the recording. They were estimated by fitting each unit’s spike amplitude distribution with a Gaussian function^47,48^ and quantifying the fraction of area under the curve clipped at the noise threshold. A 30-second segment was deemed acceptable if it had less than 5% of false positive rate <u>and</u> less than 5% of false negative rate. Acceptable intervals were concatenated and used for subsequent classifier training. A unit was required to have 3 minutes of acceptable isolation during the baseline period to be included in the sample.
● **Monkey**: Following each recording session, individual action potentials were assigned to putative neural units using the semi-automated “Full Binary Pursuit” sorter^122^, designed to distinguish temporally and spatially overlapping spikes from different neurons. Following automated sorting, we manually curated our dataset, removing neurons with significant interspike interval violations or low signal-to-noise ratios. The majority of units in our primate dataset significantly exceeded the metrics used for automated curation of the mouse data, which potentially biases our sample of primate units towards those that are easier to record.

#### Data harmonization

To achieve consistency of data acquired across labs and setups, we implemented several procedures (Supplementary Figure 1):

1. We reprocessed the wideband voltage recordings from monkeys and mice where the SpikeGLX acquisition filter was off with a causal first-order Butterworth high-pass filter (300 Hz cutoff) to agree with the hardware filter used by Neuropixels probes. Following filtering, we used a drift- and shift-matching algorithm to generate mean waveforms for each recorded unit.
2. We sought to remove one source of waveform variability by flipping the waveform, if necessary, so that the largest peak was always negative. We did so with the knowledge that the polarity of the action potential waveform depends on a number of factors including the proximity of the recording electrode to the dendrites, soma, and axon^94,123^ and relative orientation of the recording contact and the reference.
3. We preprocessed all waveform templates by selecting the mean waveform from the highest amplitude channel, resampling it to 30 kHz (if necessary), aligning it to the peak, and inverting it if necessary (see #2 above) to ensure the most prominent peak in the waveform was always negative. We used the harmonized waveforms to compute summary statistics (Figure 5, Supp Figure 4), which have been previously used to classify cerebellar neurons^19,20,62,63^.
4. We sub-sampled the spikes of each neuron by grouping waveforms with a similar amplitude on the principal channel, and therefore the same drift-state (i.e. position of probe relative to the recorded neuron): “drift-matching”.
5. We re-aligned the spikes in time by maximizing the cross-correlation of each spike to a high amplitude template: “shift-matching”. After alignment, the individual spikes were averaged, resulting in the final mean waveform for the neuron under study. Neuropixels data processing (non-manual curation, filtering, drift-shift-matching) was performed using the NeuroPyxels library^124^.

### Identification of units directly responsive to optogenetic stimulation

Units recorded during optogenetic activation experiments were deemed to be directly responsive to photostimulation if they met the following conditions: (1) their firing rate increased (ChR2) or decreased (GtACR2) more than 3.3 standard deviations from the pre-stimulus baseline within 10 ms of stimulation onset in the ’post-blocker’ trials (computed using 0.1 ms bins smoothed with a causal Gaussian filter with a standard deviation of 0.5 ms), (2) they were recorded at a depth at which pharmacological blockade was confirmed, and (3) the spike waveforms evoked in the ’post-blocker’ optogenetic stimulation trials matched those recorded during the pre-stimulation ’baseline’ period.

### Construction of 3D autocorrelograms

All recordings were performed in awake animals that were either head-fixed but otherwise free to move on a wheel (mice) or performing discrete trials of smooth pursuit (primates), so that firing rates modulated across the experimental session. To account for the impact of changes in firing rate on measures of firing statistics, we constructed “three-dimensional autocorrelograms” (3D-ACGs). At each point in time, we estimated the instantaneous firing rate of the neuron as the inverse interspike interval^125^. We smoothed firing rates using a boxcar filter (250 ms width) and evaluated the smoothed firing rate at each spike. Finally, we determined the distribution of firing rates from all interspike intervals in a recording, stratified firing rate into 10 deciles, and computed separate 2D-ACGs for the spikes in each decile. We visualized the resulting 3D-ACGs as a surface where the color axis corresponds to the probability of firing, the y-axis stratifies the firing rate deciles so that each 3D-ACG contains 10 rows, and the x-axis represents time from the trigger spike. Note that the spike counts in the autocorrelograms have been divided by the width of the bin so that the y-axis or color map is calibrated in spikes/s.

As input to the classifier, we used log distributed bins relative to *t=0* in contrast to the linearly spaced bins shown in the Figures and Supplemental material.

### Human expert labeling of cerebellar units

● **Mouse**. We used Phyllum to identify the layer of each recording. Most Purkinje cells were identified by the presence of both simple spikes and complex spikes and complex-spike-triggered histograms that showed a characteristic pause in the simple spike firing rate following the complex spike. We identified a number of recordings as Purkinje cells by the presence of simple spikes without a complex spike, location in a Purkinje cell layer, and regular firing rate resulting in characteristic “shoulders” present in the autocorrelogram. Putative molecular layer interneurons were identified by their presence in a molecular layer with firing rates above 5 spikes/s, incompatible with the firing properties of the dendritic complex spikes. A subset of putative molecular layer interneurons yielded a properly timed spike-triggered inhibition of an identified Purkinje cell simple spike. Putative mossy fibers were in a granular cell layer and some displayed a characteristic triphasic shape due to the negative afterwave recorded near the glomerulus^50,51^. Putative Golgi cells were in the granular cell layer and had broad waveforms and relatively regular firing rates. In addition, some pairs of putative Golgi cells showed a double peak in the millisecond range in their cross-correlograms, indicative of gap-junction coupling^126^.
● **Monkey.** We classified recordings as ground-truth Purkinje cells if they demonstrated the characteristic post-complex-spike pause in simple-spike firing. Units that exhibited known characteristics of Purkinje cell simple spikes but lacked a complex spike were treated as “putative” Purkinje cells and used in the comparison of classifier-predicted and expert-predicted labels. We included molecular layer interneurons only if they showed spike-triggered inhibition of an identified Purkinje cell’s simple spikes at short latency, leaving some potential molecular layer interneurons out of our sample. We included units as putative mossy fibers only if the waveform showed a negative after-wave, characteristic of recording near a single glomerulus^50,51^. We note that our classification of mossy fibers is highly conservative and likely leaves a large subset of mossy fiber recordings not near a glomerulus as unlabeled. Putative Golgi cells were identified by their presence in the granule cell layer, broad waveforms, and highly regular firing, consistent with previous recordings^63^. Expert labeling of units in the monkey were performed before collection and analysis of the ground-truth units in the mouse.

### Cross-validated cell-type classification

We began the design of our cell-type classifier by selecting the feature space passed to the model, the model class, and model characteristics such as number of units and learning rate, collectively the features that define the model’s “hyperparameters”. Our decision to select hyperparameters independently from the ground truth dataset was critical to ensure generalizability by minimizing overfitting^127^. To construct an unbiased feature space to train the model, we decided *a priori* that the model’s inputs would be anatomical location, extracellular waveform, and firing statistics. We elected not to use summary statistics because they provide an impoverished set of information compared to the inputs we selected. We optimized the model’s architecture fully independently from our ground truth dataset by leveraging n=3,090 curated but unlabeled units that were recorded in the experiments used to create the ground-truth library but were not activated optogenetically. We trained variational autoencoders to reconstruct the waveforms and 3D-ACGs of the unlabeled units, and optimized the architecture of the autoencoders based on the quality of the reconstruction, independently from the ground truth dataset. In the final classifier, we used encoder networks of the two autoencoders to reduce the dimensionality of the waveforms and 3D-ACGs of the ground-truth library. The output of the autoencoders, along with the layer of each neuron, served as inputs used to train the final classifier on the ground truth dataset. Thus, no aspect of the model’s feature space or architecture was chosen based on the model’s performance on the ground truth dataset.

Our classifier is a “semi-supervised” model because the variational autoencoders were tuned and trained with unsupervised learning on a set of unlabeled neurons while the complete classifier was trained with supervised learning on a separate set of ground-truth identified neurons. We derived our strategy from the “M1” model^128^.

### Variational autoencoder pre-training on unlabeled data

We trained two separate autoencoders to reconstruct the waveforms and log-scaled 3D-ACGs of our unlabeled units. Ultimately, the encoder networks of the autoencoders, trained on our unlabeled data, compressed the input data into two 10-dimensional ‘latent spaces’ for the 2 input features: 3D-ACG and waveform. The training objective of the variational autoencoders was the ‘Evidence Lower Bound’ loss^70^ modified to include a β term to encourage disentanglement of the latent space (Higgins et al. 2016). During training, we employed a Kullback–Leibler divergence annealing procedure to enhance model stability and convergence^129^. Both variational autoencoders were trained through gradient descent with the Adam optimizer, complemented by a cosine-annealing learning rate strategy with periodic warm restarts^130^.

To both facilitate model convergence and yield high-quality reconstructions, we manually adjusted variational autoencoder parameters to adapt the model to our specific data characteristics and improve its performance in subsequent tasks. This procedure did not rely on the ground truth dataset, so we could adjust hyperparameters freely without overfitting to the classification task.

● The final *waveform variational autoencoder* consisted of a 2-layer multilayer perceptron (MLP) encoder with Gaussian Error Linear Units (GeLU) non-linearities^131^ and a 2-layer MLP decoder also with GeLU non-linearities. It was trained for 60 epochs with η=1e-4, β=5 and a mini-batch size of 128.
● The final *3D-ACG variational autoencoder* consisted of a 2-layer convolutional neural network (CNN) encoder with average pooling after convolutions, batch normalization, and rectified linear unit (ReLU) non-linearities, and a 2-layer MLP decoder with ReLU non-linearities. It was trained for 60 epochs with η=5e-4, β=5 and a mini-batch size of 32.

The analysis described in Supplementary Figure 7 ensures that our trained variational autoencoders accurately captured the variance in our data.

### Semi-supervised classifier

The final classifier model consisted serially of: (1) the waveform and temporal feature “variational autoencoders” pretrained on unlabeled data to reduce the dimensionality of the input features; (2) a multi-headed input layer that accepted the latent spaces of the waveform and 3D-ACG variational autoencoders, along with a “one-hot-encoded” 3-bit binary code of the unit’s cerebellar layer; (3) a single fully-connected hidden layer with 100 units that processed the 3 normalized inputs; (4) an output layer with one output unit for each of the 5 cell types (Figure 6C). The value of the output units sums to 1 via a softmax function so that the output of the classifier is the probability that a given set of inputs is from each of the 5 cell types. Between the input (2) and fully-connected (3) steps, we applied batch normalization^93^ to equate the contributions of waveform, discharge statistics, and layer. The fully-connected hidden layer had a Gaussian prior that encouraged each network unit to have activation values across the training set with zero mean and unit variance. We trained the weights of the complete classifier on the data in the ground-truth library using gradient descent with a leave-one-out cross-validation strategy. We trained the models until convergence or for 20 epochs, whichever came first, with η=1e-3, a mini-batch size of 128 and the AdamW optimizer^130^. We allowed the weights in the pre-trained variational autoencoders to change during optimization to allow fine-tuning that caused a small improvement in performance on the downstream classification task.

Finally, we took several steps to ensure that our models are robustly trained and capable of generalizing well across datasets:

1. To account for “class imbalance” created by the different number of neurons in each cell type, we performed random oversampling of the under-represented cell types for every model after splitting into testing and validation data^132^.
2. We assessed the performance of all models through leave-one-out cross-validation, which has a lower bias and comparable variance to other cross-validation methods^133,134^ and has been used in the past to model small datasets such as ours^20^. Thus, leave-one-out cross-validation is better than other cross-validation methods but worse at estimating the generalization error than having an independent test set. We note that our models seemed to perform well on independent “expert-classified” test sets.
3. We did not tune the hyperparameters of the final semi-supervised classifier or of its training procedure. We used only predefined heuristic values and trained until convergence.
4. We adopted a strategy to prevent *confidence miscalibration*, the tendency of deep neural networks to exhibit over-confidence in their predictions^135^. We corrected the overconfidence of each model instance by applying a last-layer Laplace approximation to the output layer^89,90^. Further, for each leave-one-out sample, we created a “deep ensemble”^136^ by training an ensemble of 10 models with random initial conditions. We then averaged the probability for each cell type across model instantiations. Each model generated an average prediction probability for each cell type, yielding a set of 5 values (for the 5 classes) that summed to 1. To quantify classifier confidence, we averaged the predicted probability for each cell type across the 10 instantiations of the model and computed the *confidence ratio* as the ratio of the highest-to second-highest predicted cell-type for the input features from each cell in our samples. We chose a confidence ratio of 2 as the *confidence threshold* here, but higher thresholds could be applied to increase confidence in each prediction of cell type.

### Generalization of prediction to unlabeled mouse and macaque cerebellar neuron cell type

We predicted the cell type of mouse (*Medina*) and macaque (*Lisberger*) cerebellar neurons that were not involved in the classifier training procedures using an ensemble classifier that utilized all ground truth neurons and initial conditions (202 x 10 = 2020 models in total). Each of the 2020 models was slightly different from the others due to the combination of the 10 initial conditions and the ’leave-one-out’ procedure used to train them and produced slightly different results. The predicted cell-type of each neuron in the unlabeled sample was chosen as that with the maximum average prediction across the 2020 models. We applied the *confidence ratio* and *confidence threshold* as we had for the ground-truth library.

### Evaluation of differential activity patterns across cell types during behavior

We leveraged our classifier to investigate the temporal profiles of distinct populations during of classifier-labeled neurons in four behavioral paradigms during discrete trials:

● *Häusser:* Data were recorded in mice performing self-initiated locomotion. Five days after surgery mice began water restriction and habituation to head fixation on the running wheel. Mice were head fixed on the wheel once daily for about 30 min and were rewarded with drops of water for moving forward on the wheel. On the day of the recording, mice typically ran 150m in about 1h. Locomotion was recorded with a Vision Mako U-130B camera at 100 frames/s and the ipsilateral forepaw of the animals was tracked with DeepLabCut^137^. Forepaw swing onsets were identified by thresholding the paw trajectory in polar coordinates (phase obtained from Hilbert transform) and used to align neural data.
● *Hull:* Mice were water restricted for at least five days prior to experiments and then habituated to head fixation on a freely moving wheel. A tube for reward delivery was placed in front of the mouse with an IR LED and photodiode positioned to detect licks. Reward consisted of water sweetened with saccharin (10 mM), delivered every 23-85 seconds via a solenoid, for a total of 218 trials. We aligned the simultaneously-recorded responses of neurons in a single session on the time of reward delivery.
● *Medina:* Mice were trained using a classical conditioning experiment in which the presence of an LED (conditioned stimulus) predicted the occurrence of a puff to the cornea (unconditioned stimulus) 220 ms later. The exact experimental protocol has been extensively described previously^138^. Animals had been extensively conditioned and were generating reliable conditioned responses before neural recordings took place. We aligned the simultaneously-recorded responses of many neurons from a single session on the onset of the conditioned stimulus.
● *Lisberger:* We recorded neural data during discrete trials of smooth-pursuit target motion^119^. Animals were seated in front of the CRT monitor and trained to pursue the smooth movement of a black dot as it moved in one of eight directions at a constant velocity. Here, we included only trials in which the target moved in the horizontal direction towards the side of the cerebellum where we were performing neural recordings and where the monkey successfully tracked the target and maintained fixation after the termination of target motion. Data were aligned to the onset of a 650 ms duration target motion. Unlike the presentation of data from a single recording session in the experiments on mice above, we constructed a pseudo-population of neurons during smooth pursuit across *n=163* behavioral sessions across three monkeys, consistent with previous analyses of population dynamics in monkeys^72,74^.

### Neural data trajectory analysis

We included in our analysis each recorded neuron that our classifier was able to label with a confidence ratio > 2, analyzing only a single session for the 3 behaviors in mice and a pseudo-population recorded across sessions for the 1 behavior in monkeys. We converted spike trains into firing rates and temporally smoothed them using kernels that were appropriate for each behavioral paradigm. The differences in time-scales of the behavioral tasks (e.g., approximately 200 ms during reward conditioning but longer than 1,000 ms during smooth pursuit) necessitated the use of task-specific smoothing methods. We note, however, that our general conclusions are robust to a large range of smoothing time constants. Following smoothing, we normalized the firing of each neuron by the standard deviation of its activity during a 100 ms pre-trial baseline period, averaged the responses to form peri-stimulus time histograms for each neuron and subtracted the mean firing rate during the pre-trial baseline period. Changes in the normalized firing rate are therefore expressed relative to their baseline activity in units of baseline firing rate standard deviation. The PSTHs in Figure 9B are averaged across all instances of each cell type and the head maps in Figure 9C show the individual PSTHs for each cell type in the population.

A more sophisticated approach for studying the temporal structure of large-scale population responses relies on dimensionality reduction techniques to understand consistent temporal signatures that exist across neurons in a population^72–74,76,139^. To answer whether low-dimensional representations of population activity from specific cell types were different from those of a cell-type agnostic population, we derived an analysis pipeline where we could compare the neural trajectories from different sized populations in a common space. In each case, we began by constructing a matrix *X* with dimensions *NxT* where each row contained the baseline normalized and smoothed peri-stimulus time histogram with *T* time points for one of *N* neurons. To apply principal component analysis, we centered the firing rates in each row of *X*. As principal component analysis identifies the dimensions in order of variance, neurons that show dramatic differences relative to their baseline firing rate will likely drive a majority of the variance. To mitigate both the over-representation of highly responsive neurons in the absence of normalization as well as the over-representation of non-responsive neurons in the case of z-scoring, we took advantage of a previously described approach for firing rate preprocessing using a “soft” normalization procedure^72^. We reduced the influence of highly active neurons, those with modulation that exceeded their baseline standard deviation by more than 2x, by ensuring that their range was approximately unity. The range of firing rate changes for non-responsive neurons, in contrast, were reduced to values near zero and, therefore, would not contribute meaningfully to the largest principal components.

We decomposed our data matrix 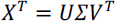 using singular value decomposition. Here, *U* represents the left singular vectors of *X^T^*, consisting of an orthonormal set of temporal representations of length *T* (principal axes across time) ordered by their variance. Typically, principal component analysis projects the data matrix *X* into a lower dimensional subspace by selecting the first *C* axes of *U*, weighted by their associated singular values contained in the diagonal elements of *Σ*, namely 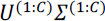. While each directional axis defined in *U* has unit magnitude, the resultant projection is scaled by the singular values whose values are related to the standard deviation, σ, of the discovered principal axes and thus have a dependence on the population size, *N*. To account for the dependence of the projection into principal component space on the population size, we transformed the singular value matrix as 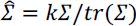. In this equation, *k* is an arbitrary constant (chosen to be 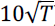) and *tr* denotes the trace operator. We can now project *X* into principal component space in a manner that retains the relative scaling between the projected axes but remains agnostic to the underlying population size using 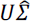. Given this formulation, we could compute the projection of each population into their respective principal component spaces even though the population sizes differed. Once we obtained our projection 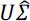, we subtracted off the mean of the activity in each principal component during the pre-trial baseline period. Therefore, all trajectories are guaranteed to start near the origin.

We next asked whether the low-dimensional geometry of each cell-type population demonstrated comparable trajectories to the label-agnostic population. To account for the fact that the sign of each principal component projection is arbitrary, we optimally rotated and reflected the low-dimensional trajectory representations of each cell-type population to align its trajectory to the low-dimensional projection of the full population, agnostic to cell-type. We did so by solving the orthogonal Procrustes problem, resulting in an orthogonal matrix *R* that reflected and rotated cell-type population trajectories about the origin to maximally agree with the cell-type agnostic trajectory, thereby finding the *R* that minimized 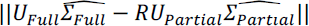. After computing *R* for each population, we analyzed the Euclidean distance between the optimally rotated cell-type trajectories and the cell-type agnostic population trajectories in the same low-dimensional space.

Because the population sizes were unequal across cell-types, we used permutation testing to assess statistical significance of the distance between optimally rotated and reflected trajectories of each cell type and to the cell-type agnostic trajectories derived from the overall population. We tested the null hypothesis that random selection of equal numbers of cells for each of the compared trajectories would show a similar distribution of Euclidean distances. We performed 1,000 permutations where we sampled neurons from both compared populations, optimally rotated and reflected these permuted trajectories into the same space, and then derived the null distribution of distances. From the empirically derived null distribution we could directly assay the significance of distances between two population trajectories.

We also tested the performance of the trajectory analysis when we introduced random errors in the cell-type labels provided by the classifier. For each given fraction of cells relabeled, we performed 1,000 replicates of randomly selected sets of cells to receive a new cell-type label from labels corresponding to either a Purkinje cell simple spike, Golgi cell, mossy fiber, or a molecular layer interneuron, with equal probability. We then selected all neurons with a given cell-type label, performed dimensionality reduction as described above, and computed the distance between the optimally rotated and reflected low-dimensional trajectory and the trajectory agnostic to cell type labels. We used bootstrapped statistics to derive the 95% confidence intervals as a function of the fraction of cells that were randomly assigned a new cell-type label.

## Author contributions

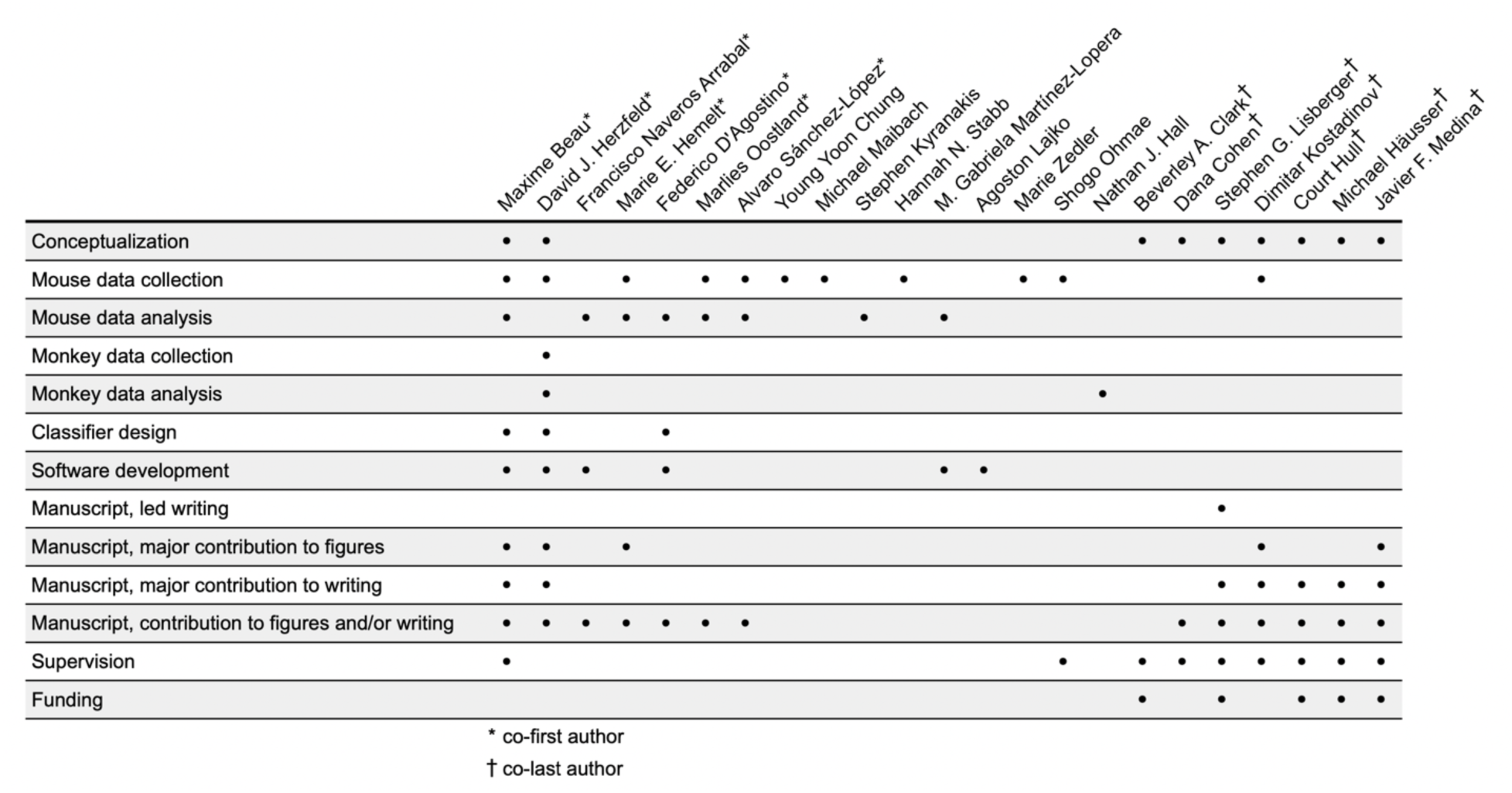

## Acknowledgements

We thank Bonnie Bowell, Soyon Chun, Wenjuan Kong, Caroline Reuter, Margaret Conde Paredes, and Stefanie Tokiyama for technical support; and Arnd Roth for helpful discussions. Funding provided by: NIH grants R01-NS112917 (SGL, JFM, CH), K99-EY030528 (DJH), R01-NS092623 (SGL), R01-MH093727 (JFM); ERC AdG 695709 (MH); Wellcome Trust PRF 201225/Z/16/Z and 224668/Z/21/Z (MH) and 225951/Z/22/Z (DK); EMBO ALTF 914-2015 (DK); European Union’s Horizon 2020 research and innovation programme under the Marie Skłodowska-Curie grant agreement No 844318 (MO) and No 891774 (FN); SYNCH project funded by the European Commission under the H2020 FET Proactive program-Grant agreement ID 824162 (DC)

## Data availability

The data that support the findings of our study will be made publicly available at the time of publication.

## Code availability

The custom analysis code used in our study will be publicly available on Github at the time of publication for major packages. Other code will be available from the corresponding author upon request.

