## Supplemental Material for "A deep-learning strategy to identify cell types across species from high-density extracellular recordings"

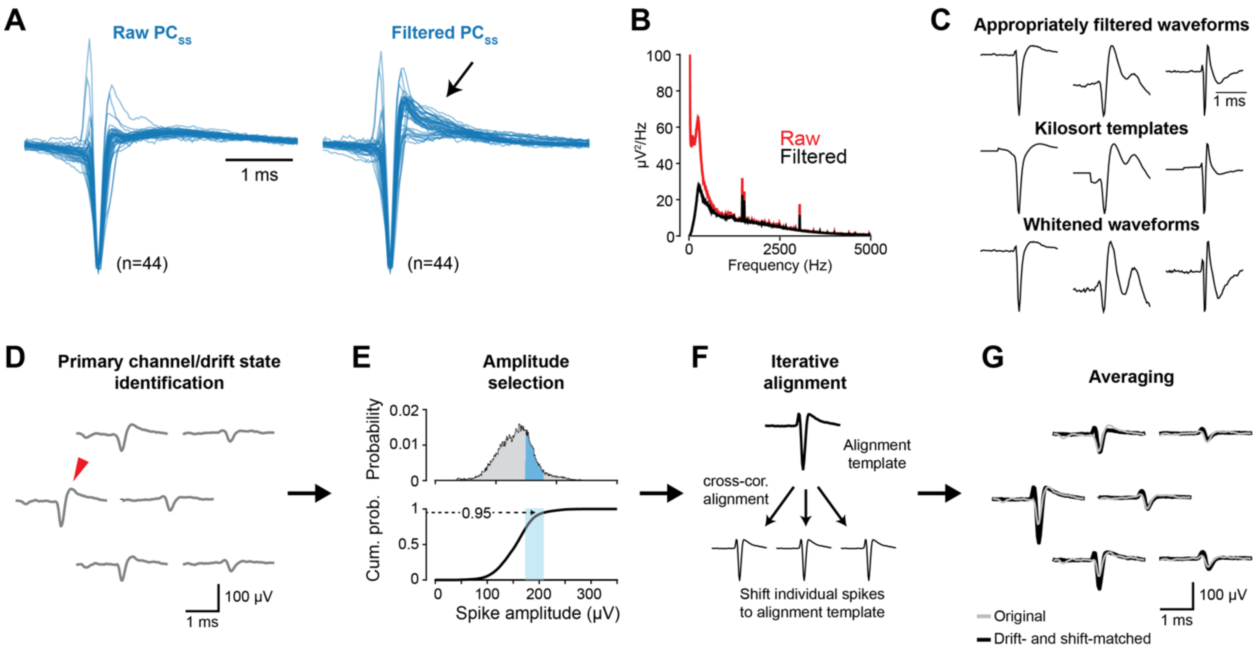


**Supplementary Figure 1: Analysis pipeline for preprocessing of neural waveforms.** As we compared recordings across labs and understood how the idiosyncrasies of different analysis pipelines work, we learned that we needed to harmonize data across preparations and to ensure that the output of our pipeline provided the best possible estimate of the actual waveform. Careful preprocessing of the waveform reduces the variation in waveform within each cell type and probably improves the performance of the classifier at distinguishing cell types. We detail those procedures - consistent filtering and drift-shift matching - here.

**A.** Simple-spike waveforms from 44 ground-truth Purkinje cells, identified through their complex-spike-induced pause in simple-spike firing. Traces on the left display normalized voltage-versus-time traces from the primary channel of Neuropixels recordings with the hardware filter disabled. Traces on the right depict the same waveforms after application of a software filter that is equivalent to the onboard hardware filter on Neuropixels probes (single-pole 300 Hz high-pass Butterworth). The black arrow points out the main difference in waveform shapes before versus after filtering. The procedure to harmonize waveforms across datasets by applying a software version of the hardware filter was critical. **B.** Representation of power in the low-frequency band averaged across the primary channels for raw (red) and filtered (black) waveforms from panel A, verifying the impact of the onboard hardware filter. **C.** Comparison of waveforms for three cerebellar neurons, showing how the standard tools used with Neuropixels probes can cause aberrations in the waveforms: top row shows the best reconstruction of waveforms of 3 units, using the analysis pipeline developed in our study; middle row shows the templates created by Kilosort, which can be quite distorted relative to the best waveform identified by our process; bottom row shows the results of zero phase component analysis (ZCA) whitening by Kilosort, a process that is relevant to performing spike-sorting but that can badly distort the waveform because of the role of activity on neighboring channels in ZCA whitening. The distortions of some waveforms by the standard analysis pipeline underscores the improvements we have made to provide classifier inputs of the highest quality. **D-G.** Preprocessing pipeline for high-quality neural waveform identification using the drift-shift-matching algorithm. **D.** Z-drift matching: identification of the primary channel through the largest peak-to-peak amplitude (red arrow). **E.** X-Y drift-matching: sub-selection of neural waveforms on the identified primary channel, with the top plot showing the distribution of peak-to-peak amplitudes and the bottom plot depicting the cumulative probability distribution. A subset of N (user-configurable) action potential waveforms with peak-to-peak amplitudes below the 95th percentile is selected (blue shaded region), eliminating spikes in the 95th to 100th percentiles to mitigate potential large amplitude artifacts. **F.** Shift-matching: consecutive/iterative alignment of small batches to waveforms via the peak in cross-correlation to an alignment template, computed as the mean of the largest peak-to-peak waveforms following amplitude selection. **G.** Final averages of the aligned sub-batches of waveforms. The black curves depict the waveform template following the complete drift-shift pipeline, while the gray waveforms show the original mean waveform reproduced from (D). The difference between the gray and black waveforms shows the impact of our analysis pipeline.

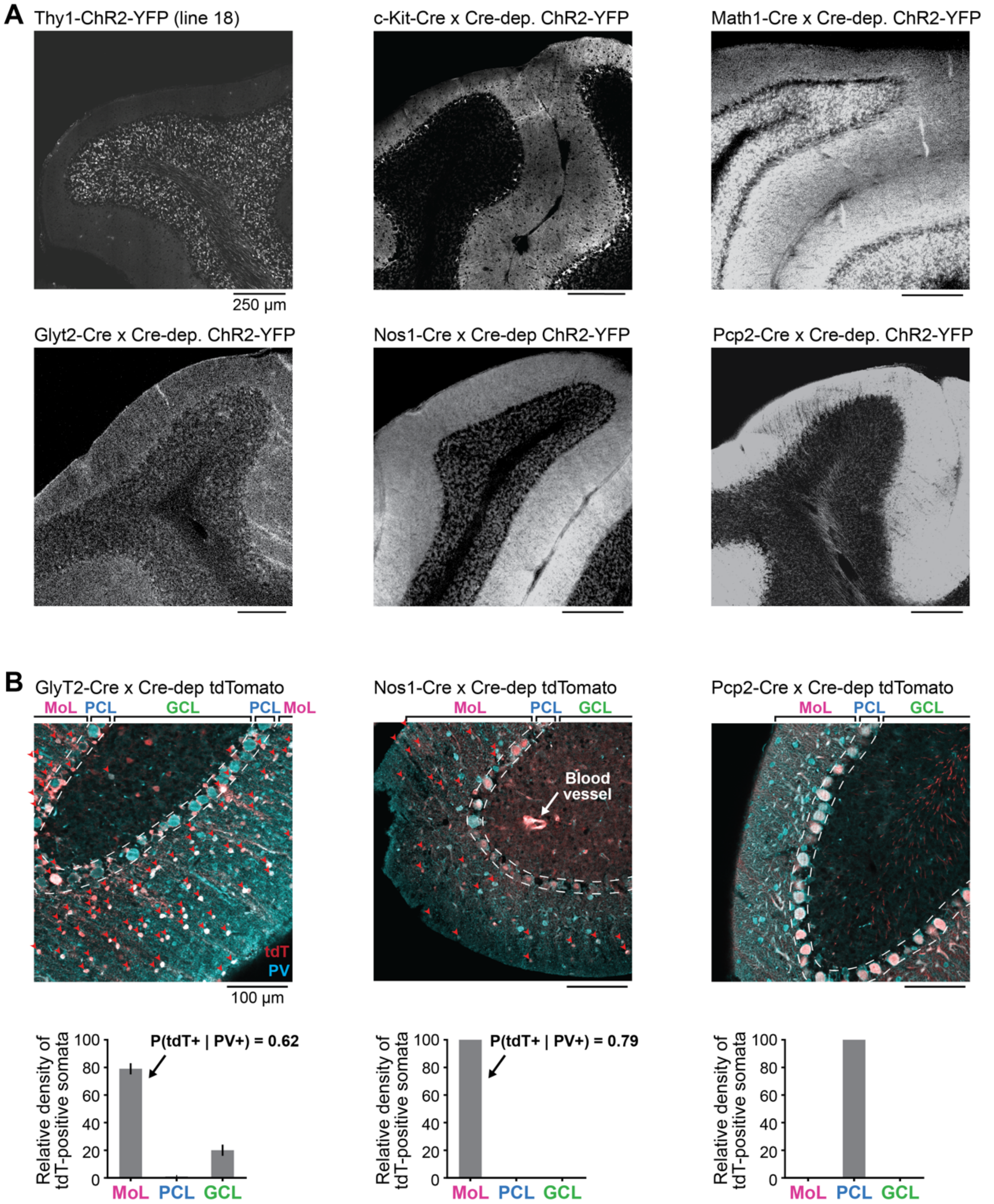


**Supplementary Figure 2: Quantifying the presence or absence of off-target expression of opsins in the mouse lines used in our study**. Off-target labeling is an important feature in some of the Mouse lines we used. As a consequence, we could not simply assume that an optogenetically-activated neuron was of the cell-type that a line has previously been described to label. Instead, we developed a strategy to manage off-target labeling based on (i) careful assessment of histology for all the mouse lines we used and (ii) identification of the layer of the recordings by Phyllum. Here, we show histology and immunohistochemistry for the main lines we used.

**A**: Histology showing the localization of the ChR2-YFP fusion protein in our mouse lines. The top row shows that off-target expression was minimal in the Thy1-line-18 used to identify mossy fibers, the c-kit-Cre line used in one laboratory to identify molecular layer interneurons, and the Math1-Cre line used to attempt to identify granule cells. The second row shows that the GlyT2-Cre line used in one laboratory to identify Golgi cells has substantial off-target expression in molecular layer interneurons, the Nos1 line that we used to identify molecular layer interneurons shows non-neuronal along with dense labeling in the molecular layer, and the L7-Cre line that we used to identify some ground truth Purkinje cells shows very little off-target labeling. **B**: Co-localization of Cre-expressing and parvalbumin (PV)-expressing neurons in the GlyT2-Cre, Nos1-Cre and L7-Cre lines, using tdTomato expression in the Ai9 as a proxy for Cre. Histograms below each image show distribution of tdTomato-positive cells across cerebellar layers in each line. Notably, all tdTomato-positive cells in the molecular layer were also PV-positive, confirming that they are molecular layer interneurons.

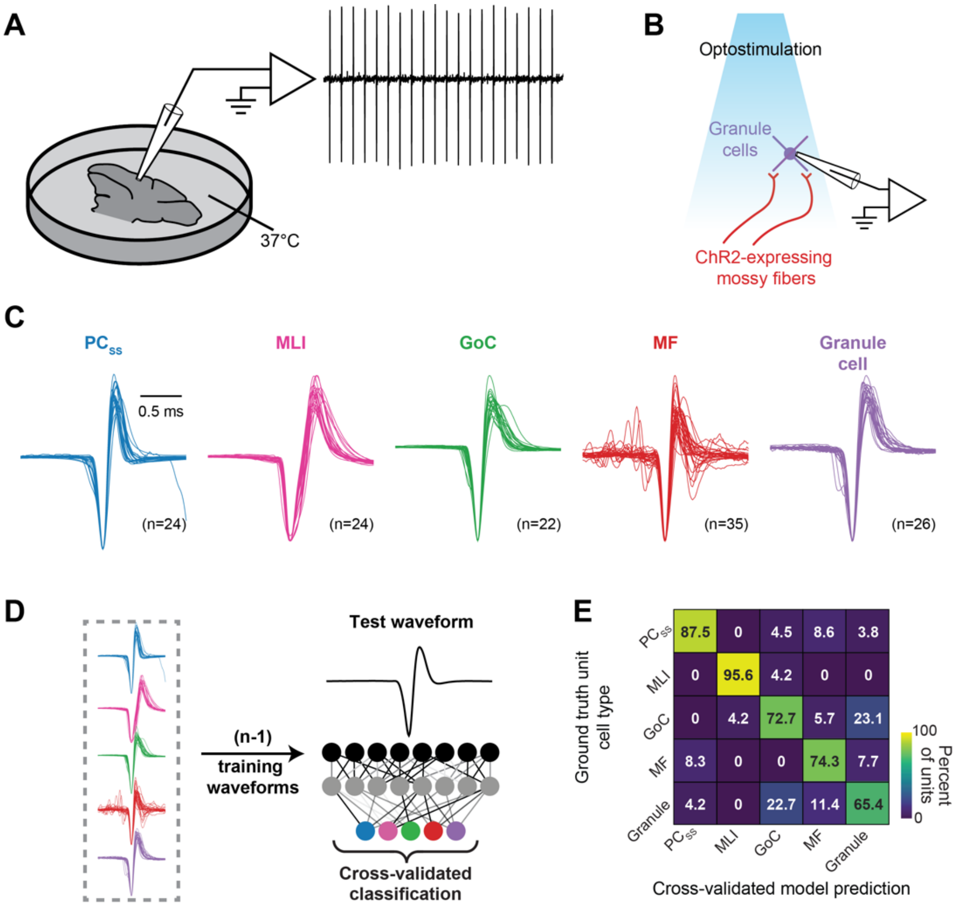


**Supplementary Figure 3: Demonstration from *in vitro* recordings that waveform is informative about cell type.** We were provoked, by the impression from Figure 5C that waveform from extracellular recordings is a useful indicator of cell type, to perform one more set of experiments to test the informativeness of waveform in a different kind of ground-truth data. We performed cell-attached patch recordings in slices of the mouse cerebellum at 37$^{o}$ C. We identified mossy fibers using optogenetics with the Thy1-ChR2 line 18 and we identified other cell types by visualizing them in the microscope during recording. The currents associated with action potentials were both very uniform within cell types and clearly different between cell types. It is not surprising that the waveforms are more uniform in the *in vitro* recordings compared to the extracellular fields recorded *in vivo* with multi-contact probes. Many uncontrolled factors affect the exact waveform recorded extracellularly. Also, the difference in recording technique and preparation precludes comparison of the waveforms recorded *in vitro* with those in Figure 5C, but the principle that different cell types have distinguishable waveforms remains. We verified the informativeness of waveform from the *in vitro* data by creating a deep-learning classifier and validating its performance with a “leave-one-out” strategy. The overall accuracy of the classifier was 78% compared to the 20% expected from random performance. We conclude that there are mechanistic physiological reasons why we can use extracellular waveform as one major feature to classify cell types.

**A**: Slice recording schematic. **B**: Schematic of optogenetic stimulation of mossy fibers during patch recording. **C:** Superimposed waveforms of identified cell types, with different colors showing different cell types: $PC_{SS}$, Purkinje cell simple spikes; MLI, molecular layer interneuron; GoC, Golgi cell; MF, mossy fiber. **D**: Schematic of a machine learning classifier that we trained to predict cell type based on waveform. **E**: Confusion matrix showing the performance of the classifier on left-out test cell types. The numbers in the entries of the matrix indicate the percentage of cells of a given ground-truth type (y-axis) as a function of the prediction of the classifier on the x-axis. The diagonal has the highest percentages, meaning that the classifier was accurate: the overall accuracy of the classifier was 78% compared to the 20% expected from random performance.

| | $PC_{SS}$ (n=69) | $PC_{CS}$ (n=58) | MLI (n=27) | GoC (n=18) | MF (n=30) |
|---|---|---|---|---|---|
| Mean firing rate (spikes/s) | 104.9 ± 38.1 | 1.2 ± 0.3 | 27.3 ± 22.5 | 18.9 ± 12.0 | 17.3 ± 15.6 |
| CV | 0.45 ± 0.17 | 0.95 ± 0.3 | 0.81 ± 0.23 | 0.76 ± 0.37 | 1.6 ± 0.9 |
| CV2 | 0.36 ± 0.09 | 0.88 ± 0.08 | 0.69 ± 0.18 | 0.6 ± 0.15 | 0.92 ± 0.28 |
| ISI std (ms) | 4.9 ± 2.6 | 905.9 ± 470.5 | 106.7 ± 183.8 | 62.4 ± 51.0 | 172.0 ± 172.8 |
| Bursting (95th percentile of inst. firing rate, spikes/s) | 201.3 ± 68.1 | 9.7 ± 1.9 | 142.3 ± 99.3 | 105.6 ± 92.2 | 507.9 ± 328.4 |
| Waveform peak-to-trough width (ms) | 0.27 ± 0.08 | 0.36 ± 0.27 | 0.42 ± 0.12 | 0.37 ± 0.09 | 0.16 ± 0.09 |
| Peak-to-trough ratio | 0.39 ± 0.17 | 0.72 ± 0.39 | 0.37 ± 0.22 | 0.61 ± 0.22 | 0.48 ± 0.26 |
| Recovery constant (ms) | 0.45 ± 0.27 | 0.4 ± 0.41 | 0.47 ± 0.2 | 0.28 ± 0.09 | 0.24 ± 0.32 |
| % spatial decay @ 24µm | 52.6 ± 15.4 | 49.7 ± 17.9 | 54.3 ± 16.9 | 40.7 ± 16.0 | 61.9 ± 15.9 |

**Supplementary Figure 4. Mean and standard deviation of commonly reported metrics used to characterize waveform and firing statistics, displayed for the cell types in the ground-truth library.** The table shows the extent of overlap across cell types for statistical measures of firing and waveform properties, to allow comparison with measures reported in previous studies[20,58,62–64,83]. Cell type abbreviations are: $PC_{SS}$, Purkinje cell simple spikes, $PC_{CS}$, Purkinje cell complex spikes; MLI, molecular layer interneurons; GoC, Golgi cells; MF, mossy fibers. For each unit, we measured metrics related to firing properties, including the mean firing rate, coefficient of variation (CV), mean CV2[140], and the standard deviation of the interspike interval distribution. We devised a metric to measure the maximal instantaneous firing rate for each unit while remaining robust to noise by computing the instantaneous firing rate as the inverse of adjacent interspike intervals and reporting the 95th percentile of the resulting distribution. We also report metrics commonly used to summarize waveform properties, including the peak-to-trough width[55], peak-to-trough ratio[54,55] and recovery constant, also called “end-slope”[54]. Finally, we computed a metric to summarize the spatial decay of each unit’s electrical footprint across channels of the Neuropixels by computing the percentage of the peak-to-peak amplitude measured on each unit’s largest channel that was lost on adjacent diagonal contacts, 24 µm away.

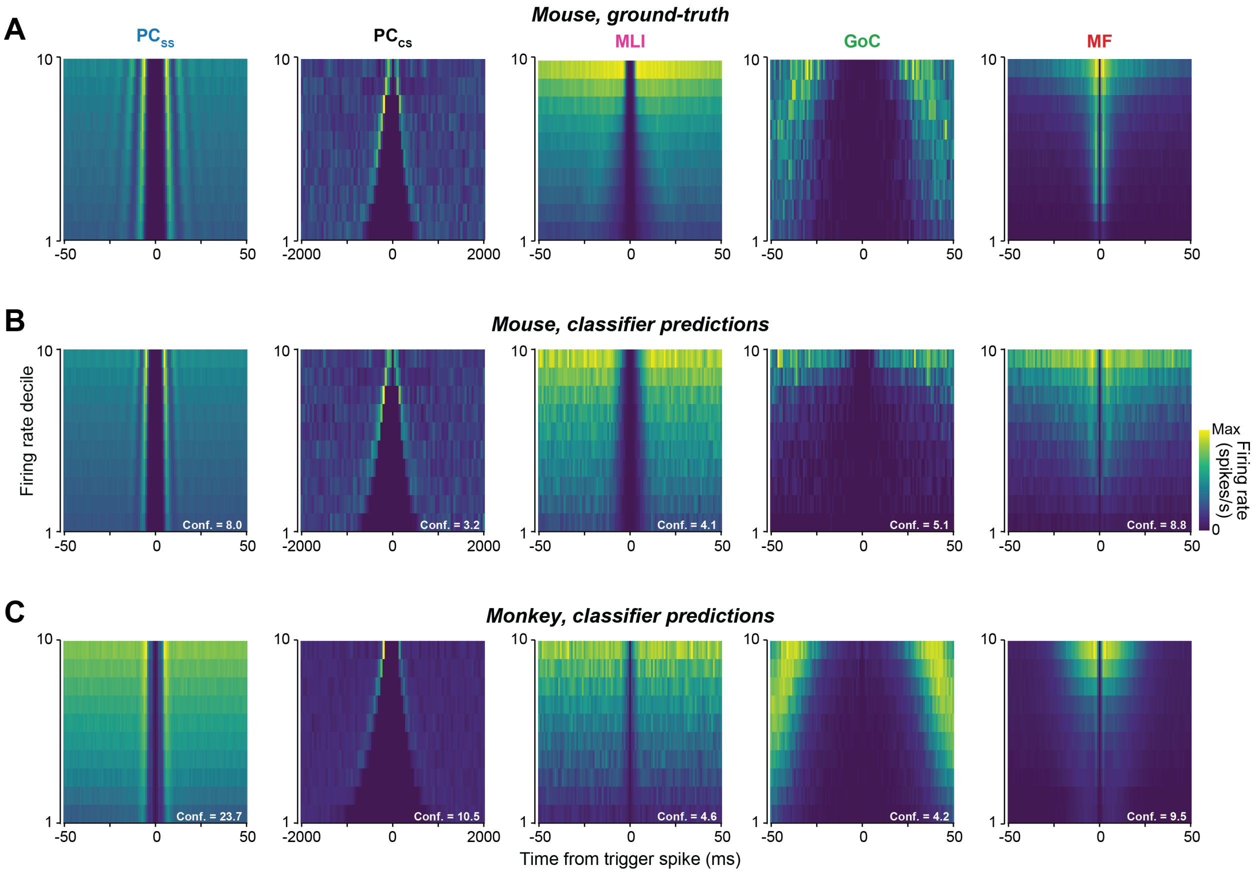


**Supplementary Figure 5. Example three-dimensional autocorrelograms for 5 cell-types in the ground-truth library from mice (A) and from non-ground-truth recordings in mice (B) and monkeys (C).** For the non-ground-truth recordings in B and C, we selected examples where the prediction of the classifier agreed with the experts' identification of cell-type. Cell types are: $PC_{SS}$, Purkinje cell simple spikes; $PC_{CS}$, Purkinje cell complex spikes; MLIs, molecular layer interneurons; GoCs, Golgi cells; MFs, mossy fibers.

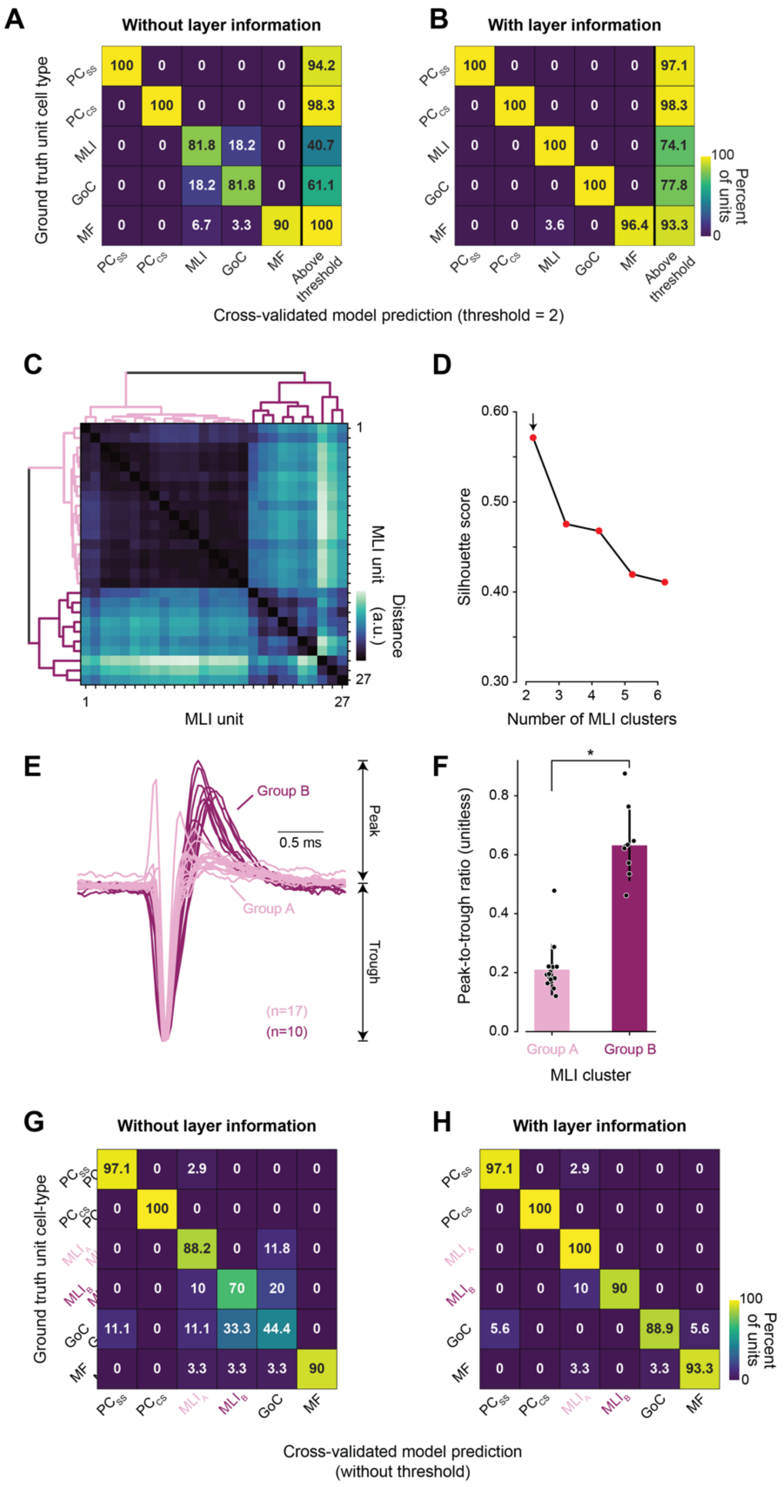


**Supplementary Figure 6: Classifier confusion without layer information as an input, and an explanation for the confusion based on the existence of two groups of molecular layer interneurons with different waveform shapes.** The classifier performs better when we include layer information as an input, especially for molecular layer interneurons and Golgi cells (A, B). When we investigated further, we realized that the waveforms of the full sample of molecular layer interneurons in the ground-truth library suggested a bimodal distribution. Here, we characterize the two groups quantitatively (C-F) and show that the classifier can distinguish between the two groups of molecular layer interneurons. Because one group of molecular layer interneurons has a waveform very similar to Golgi cells, the classifier performs better with versus without layer information as an input (G,H). The two types of waveforms in molecular layer interneurons seem likely to represent two types of interneurons but are unlikely to map onto the known types of molecular layer interneurons[71].

**A-B**. Confusion matrices showing the cross-validated performance of the classifier when layer was not (**A**) or was provided as input to the classifier (**B**). Here, the value in each entry of the matrix shows the percentage of ground-truth cell types on the y-axis that were predicted by the classifier to be the cell type on the x-axis. Note the confusion specifically between molecular layer interneuron neurons and Golgi cells without layer information (**A**). **C**. Pairwise distance matrix between normalized waveforms for the entire sample of 27 molecular layer interneurons in the ground-truth library. Lines on the left and top show dendrograms obtained via hierarchical clustering. **D**. Silhouette score as a function of molecular layer interneuron clusters. A high score indicates that a sample matches appropriately to its own cluster and is separated from neighboring clusters. Maximizing the silhouette score provides an unbiased estimate of the number of underlying clusters. Here, the assumption of two clusters maximizes the silhouette score. The maximum silhouette score is +1, minimum silhouette score is -1. **E**. Separation of the waveforms of molecular layer interneurons into group A (n=17) and group B (n=10), shown in different colors, based on the pairwise distance matrix and dendrograms in (A). **F**. Comparison of peak-to-trough ratios for the two clusters of molecular layer interneuron. **G-H**. Confusion matrices showing the cross-validated performance of the classifier with separate labels for Group A and Group B molecular layer interneurons when layer was (**G**) or was not provided as input to the classifier (**H**). Here, the value in each entry of the matrix shows the percentage of ground-truth cell types on the y-axis that were predicted by the classifier to be the cell type on the x-axis. Note the confusion specifically between molecular layer interneuron Group B neurons and Golgi cells without layer information (**G**).

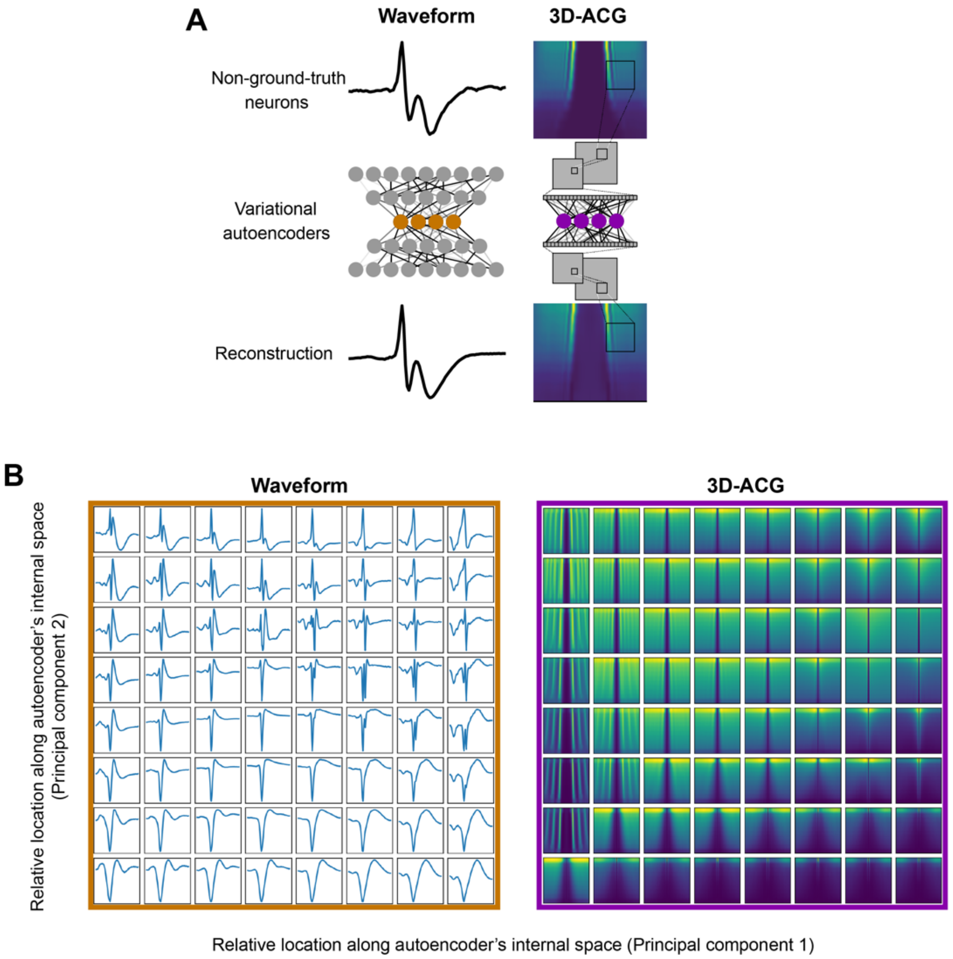


**Supplementary Figure 7. Visualization of the diversity of waveforms and 3D-ACGs captured by the autoencoders trained on mouse neurons independent from the ground truth dataset.**

We explicitly optimized the structure of the variational autoencoders to successfully reconstruct the waveform and 3D-ACG of a sample of 3090 unlabeled cerebellar neurons. However, such optimization can result in overfitting the training dataset, resulting in poor dimensionality reduction performance for our test-cases on the ground-truth and expert-labeled datasets. Therefore, we sought to qualitatively examine the extent to which each autoencoder could represent a diversity of input statistics without explicitly evaluating autoencoder performance on datasets that might later be used to test classification efficacy. Here, we show that the dimensionality reduction performed by the two variational autoencoders captures the diversity of waveform and 3D-ACG statistics present in our ground-truth and expert-labeled datasets.

A. Two variational autoencoders, one for waveform and a separate convolutional variational autoencoder for 3D-ACG, were designed to reduce the dimensionality of their respective inputs to a 10-element vector (colored circles) that could be subsequently supplied as inputs to the final classifier. Each variational autoencoder is an artificial neural network that reduces the input dimensionality by placing an information bottleneck ('latent space') between an encoding and decoding network. We trained the weights in the autoencoders with gradient descent (see

Methods) using either the waveform (left) or 3D-ACG (right) derived from a set of unlabeled cerebellar neurons. Training minimized the difference between the supplied input and the encoded-decoded output.

B. Distribution of waveforms and 3D-ACGs represented by the latent space of the autoencoders. Most of the low-dimensional representations inside the waveform variational autoencoder were occupied by variations of somatic waveforms, which are indeed the most common in the dataset. However, all other typical spike shapes were also represented, including dendritic waveforms, both bi-phasic and tri-phasic axonal waveforms, and waveforms featuring post-synaptic depolarizations. The same was true for the reconstructed 3D-ACGs, which captured activity profiles corresponding to bursting, oscillations, and both high and low firing rates. Note that some non-biological-looking traces in the figure should not be interpreted as a failure of the model but rather as a by-product of the interpolation process used to visualize the reconstructions.

To generate the graphs in panel B and evaluate whether the autoencoders could represent a broad range of input statistics in their latent space, we provided novel inputs to the decoding half of each autoencoder and visualized the resulting reconstructions[70]. Because the values supplied to the decoder could change in 10 dimensions (corresponding to the 10-dimensional latent space), to visualize the reconstructions we identified the two dimensions in the latent space that accounted for the majority of the variance across our training dataset. We utilized the encoder network to encode the waveforms and 3D Auto-Correlograms (3D-ACGs) of 3090 unlabeled units onto a latent space. Within this space, we then performed principal component analysis to identify the two components that accounted for the most variance. We generated novel inputs to the decoding half of the autoencoder via the weighted sum of these two principal components. The location of each reconstructed waveform (left) and 3D-ACG (right) in the 8x8 matrices corresponds to the relative position of the weights applied to the first (horizontal axis) and second (vertical axis) principal components. The weights were chosen as the octiles of a mean-zero Gaussian distribution whose standard deviation was chosen to yield a representative distribution of the waveforms and 3D-ACGs observed across these principal components. We chose the weights from a Gaussian distribution because the distribution of activations across each unit in the latent space was encouraged to be a zero-mean Gaussian through the assigned standard normal prior (see Methods), but other choices of weights across a similar range of values would yield qualitatively similar results. We reasoned that if the autoencoders were overfit or if the set of waveforms and 3D-ACGs in the unlabeled dataset were insufficiently diverse, we would observe discontinuities in the reconstructed outputs rather than smooth transitions that could accommodate a wide range of input statistics.
